# An RBP-J-heparan sulfate-dependent gatekeeper directs macrophage fate between inflammatory regulation and osteoclastogenesis

**DOI:** 10.64898/2026.09.02.749003

**Authors:** Ting Zheng, Huanmeng Hao, Jialin Jiang, Zhangjie Wang, Miaomiao Li, Courtney Ng, Jian Liu, Ding Xu, Baohong Zhao

## Abstract

Macrophages in bone possess a unique bi-potential capacity. Different from their primary roles in inflammation and immunity, these cells can differentiate into bone-resorbing osteoclasts. However, it remains unclear how macrophages determine which function to engage in bone, especially when exposed to the same stimulus, such as TNF, which can drive both pathways. Understanding these mechanisms is particularly important in inflammatory bone diseases, such as rheumatoid arthritis, where inflammation and bone resorption are hallmarks. Here, we identified an RBP-J–Hs2st1–2-O-sulfated heparan sulfate (HS)–IFNβ dependent molecular network that acts as a gatekeeper balancing macrophage fate inclinations in response to TNF. This network integrates cellular outside-in, inside-out, and relayed outside-in pathways. RBP-J deficiency in macrophages shifts TNF action from inflammatory regulation to osteoclastogenesis. Hs2st1, the exclusive biosynthetic enzyme for 2-O sulfation of HS, is a key target suppressed by RBP-J. Loss of Hs2st1 significantly reduces inflammatory osteoclastogenesis and arthritic bone erosion without affecting physiological bone mass. Elevated Hs2st1 expression and 2-O sulfation shift macrophages toward an osteoclastogenic fate, and vice versa. We further identified residues in IFNβ that are responsible for binding HS. Mutation of these sites diminished HS binding, augmented type I IFN response, and more strongly suppressed osteoclastogenesis. Our findings reveal a previously unrecognized molecular program fine-tuning macrophage fate and function, and highlight potential therapeutic strategies for inflammatory diseases with bone defects.

## INTRODUCTION

Macrophages, as a myeloid lineage cell type, play multiple roles in immunity and inflammation, including pathogen clearance, cytokine and chemokine production, tissue homeostasis and repair, as well as immunoregulation at various levels (*1–4*). As one of the first-line responders and multifunctional cells, macrophages are highly plastic in response to environmental cues. Recent advances have uncovered a broad spectrum of macrophage subsets beyond the traditional M1 and M2 polarization states in distinct tissue microenvironments (*4, 5*). This extraordinary plasticity allows macrophages to fulfill diverse functions under both homeostatic and pathological conditions, with broad implications for health and disease. In the skeleton, beyond the general polarization and activation seen in other tissues, macrophages possess a unique capacity to differentiate into giant multinucleated osteoclasts. These cells are the exclusive bone-resorbing cells essential for skeletal development and remodeling under physiological conditions, but also the direct drivers of pathological bone erosion in diseases (*6*).

Inflammation and bone erosion are hallmarks of multiple skeletal diseases, including rheumatoid arthritis (RA), psoriatic arthritis, and periodontitis (*6–10*). Since macrophages play important roles in both inflammation and osteoclastic bone destruction, understanding how they balance these dual roles is essential for gaining mechanistic insight and developing targeted therapies. In particular, there remains a long-standing question of how myeloid macrophages determine whether to maintain an inflammatory phenotype or commit to osteoclast differentiation in inflammatory environments, especially under the influence of cytokines, such as Tumor Necrosis Factor (TNF), which can drive both processes.

TNF is a key pathogenic cytokine in inflammatory bone diseases (*11*). It activates macrophages to produce pro-inflammatory cytokines, including IL-1, IL-6, and TNF itself. During chronic inflammation, TNF further stimulates the expression of interferon-stimulated genes (ISGs), including type I IFN response genes, such as *Mx1*, *Ifit1*, and *Ifit2*, as well as chemokines, such as *Cxcl9* and *Cxcl10*, through a trace amount of autocrine IFNβ production in these cells (*12, 13*). Through these activities, TNF stimulates macrophages to execute their primary roles in orchestrating the inflammatory response and immunoregulation. On the other hand, TNF is able to stimulate osteoclast formation and bone resorption (*14, 15*). While canonical osteoclast differentiation, which is essential for physiological bone remodeling, is primarily induced by the receptor activator of nuclear factor-kappa-B ligand (RANKL)-RANK signaling, a non-canonical, RANKL-independent osteoclast differentiation pathway has recently been identified under inflammatory conditions, driven by transforming growth factor β (TGFβ) priming and TNF stimulation of macrophages (*16*). This TNF-mediated non-canonical osteoclastogenic program is also observed in RA (*16*). However, the mechanisms by which TNF switches the balance of its pro-inflammatory and osteoclastogenic activities remain poorly understood.

Effective treatment of inflammatory bone diseases, such as RA, requires controlling both inflammation and pathological bone erosion. TNF inhibitors have demonstrated therapeutic efficacy but carry long-term immunosuppressive risks, including opportunistic infections and reactivation of latent tuberculosis (*11*). Similarly, RANKL inhibitors can limit bone loss but also disrupt physiological bone remodeling and repair, raising concerns about complications, such as atypical fractures and osteonecrosis (*17–19*). These challenges underscore an unmet clinical need to discover regulatory mechanisms that can selectively target pathological osteoclastogenesis and excessive inflammation while preserving immune function and normal bone turnover. However, such mechanisms remain largely uncharacterized and represent a critical gap in our understanding of disease pathogenesis and therapeutic strategies.

In this study, we identify an RBP-J-regulated, heparan sulfate (HS)-dependent regulatory network that serves as a gatekeeper balancing macrophage fate decisions in response to TNF. HS, a linear sulfated polysaccharide expressed on the cell surface and extracellular matrix, regulates cellular activities through interactions with a wide range of binding proteins (*20–23*). Hs2st1 is the exclusive biosynthetic enzyme responsible for adding sulfate groups to the 2-O-position of the HS chain on the cell surface (*20–23*). This 2-O sulfation modulates the interaction between HS and its binding proteins, thereby significantly influencing diverse cellular functions (*24–26*). We uncovered a unique, relayed signaling network comprising newly discovered outside-in (TNF → RBP-J, which suppresses Hs2st1), inside-out (Hs2st1 → HS 2-O sulfation → IFNβ), and relayed outside-in (IFNβ → immune function and osteoclastic inhibition) pathways that specifically regulate TNF-mediated macrophage responses and inflammatory osteoclastogenesis. In this context, the 2-O sulfated HS on the cell surface binds TNF-induced IFNβ. We further identified specific HS-binding residues in IFNβ and demonstrated that HS 2-O sulfation acts as a molecular gatekeeper modulating IFNβ activity to switch the balance between macrophage immune function and osteoclast differentiation in response to TNF. Importantly, this network appears not to affect physiological osteoclastogenesis or normal bone remodeling. The components and mechanisms identified here provide promising targets for developing therapeutic strategies that suppress inflammatory bone resorption and fine-tune macrophage inflammatory regulation while minimizing side effects on bone remodeling or immune function.

## RESULTS

### TNF acts through the RBP-J-Heparan sulfate (HS) axis to modulate macrophage inflammatory and osteoclastogenic programs

While the role of TNF in orchestrating inflammatory and immune regulation in macrophages is well established, its intrinsic capacity to induce macrophage differentiation into osteoclasts (osteoclastogenesis) is limited. The mechanisms by which TNF determines whether macrophages engage in immune regulation or osteoclastogenesis remain largely unknown. Building on our identification of RBP-J as a crucial suppressor of TNF-induced osteoclastogenesis (*14, 27*), we performed RNA sequencing using primary bone marrow macrophages (BMMs) from *Rbpj^ΔM^* (*Rbpj^f/f^;LysMcre,* LysMcre driven myeloid RBP-J conditional KO mice) and the wild-type control mice (*Rbpj^+/+^;LysMcre,* hereafter referred to as WT Ctrl) under TNF stimulation to dissect how TNF modulates the bipotential fate of macrophages. Although RBP-J is traditionally recognized as a central transcriptional regulator in canonical Notch signaling, emerging evidence indicates its broader involvement in pathways, such as TLR and TNF signaling (*14, 27–34*). Notably, TNF activates RBP-J (*34*), and RBPJ has been identified as a genetic susceptibility locus in RA (*35–37*), underscoring its importance in inflammatory diseases.

In WT control macrophages, TNF stimulation predominantly induced genes linked to inflammation, cytokine production, and phagocytosis, as revealed by Gene Ontology (GO) analysis of the RNA-seq data (Fig. 1A). Strikingly, RBP-J deficiency shifted this transcriptional program, enriching gene sets associated with osteoclast differentiation and bone resorption (Fig. 1A). Further analysis uncovered a significant and unexpected enrichment of heparan sulfate (HS) biosynthetic pathways in RBP-J deficient macrophages (Fig. 1B), suggesting a previously unrecognized regulatory axis.

**Figure 1.**
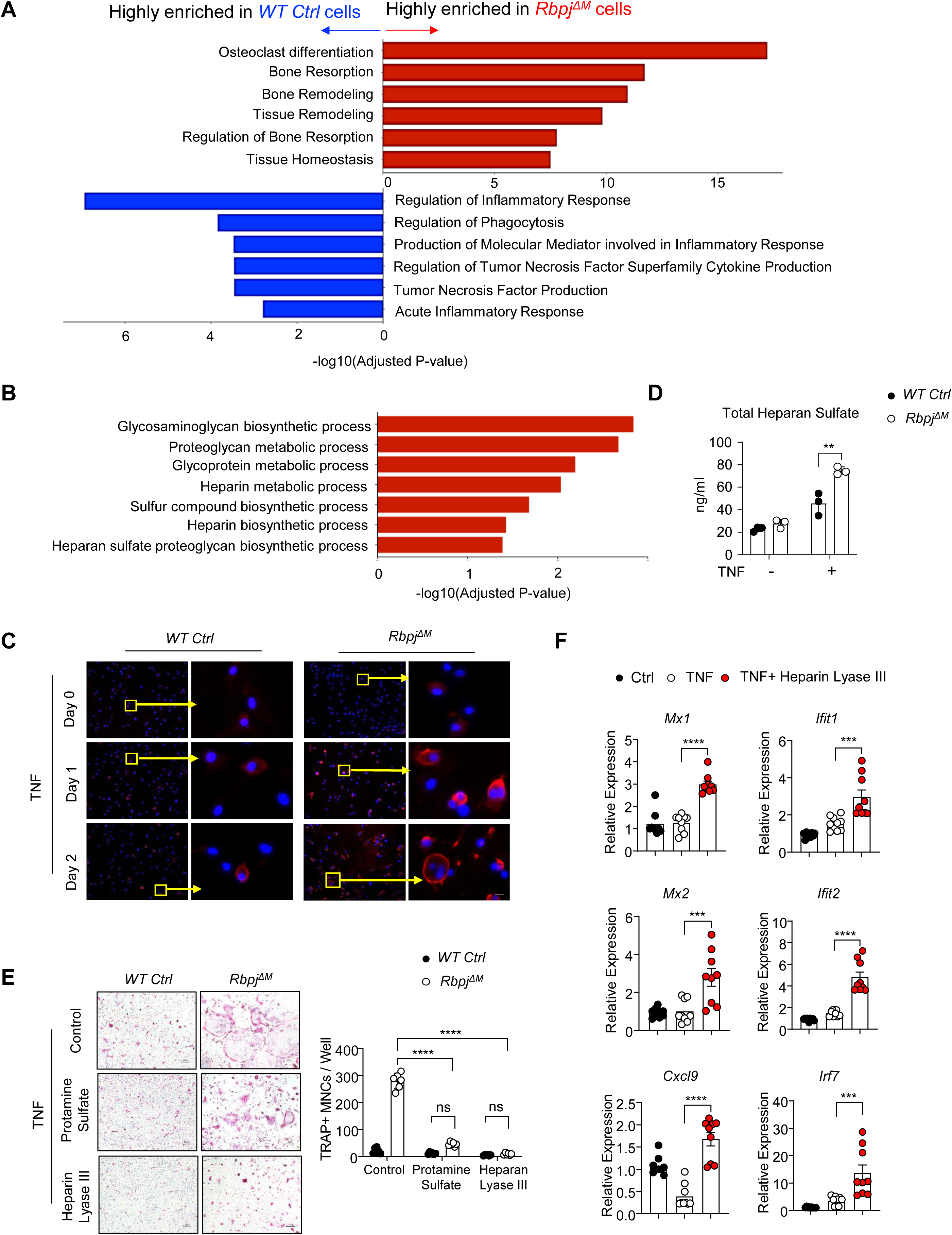
RBP-J suppresses the expression of heparan sulfate (HS), which in turn modulates RBP-J function in macrophages. (A) RNAseq based pathway analysis of differentially expressed genes (DEGs) in *WT Ctrl* and *Rbpj^ΔM^* BMMs stimulated with TNF (40 ng/ml) for 3 days. Significance is shown as –log10(adjusted p value). (B) Gene Ontology analysis of the HS biosynthetic pathways enriched in *Rbpj^ΔM^* BMMs under TNF treatment. (C) Immunofluorescence staining of HS on the cell surface. Yellow arrows: representative cells. Red: HS; Blue: Dapi as the nuclear counterstain. (D) Quantification of total HS levels in BMMs from WT and *Rbpj^ΔM^* mice treated with or without TNF for 2 days, measured by LC-MS/MS (n = 3/group). (E) Osteoclast differentiation in BMMs from *WT Ctrl* and *Rbpj^ΔM^*mice stimulated with TNF in the presence or absence of protamine sulfate (10 uM) or heparin lyase III (HL-III, 10 mIU/ml) for 3 days. Left: TRAP staining; Right: the number of TRAP-positive multinucleated cells per well (n = 6/group). TRAP-positive cells are shown in red. (F) qPCR analysis of ISG gene expression in BMMs from *Rbpj^ΔM^*mice stimulated with TNF in the presence or absence of HL-III (10 mIU/ml) for 2 days (n = 9 / group). Scale bar: C, E: 100 μm. Data are presented as mean ± SD. *p < 0.05; **p < 0.01; ***p < 0.001; ****p < 0.0001; ns, not statistically significant. Data representative of at least three independent experiments. Source data are provided as a Source Data file.

HS sulfotransferases control the biosynthesis of HS chains on the cell surface. Consistent with the GO pathway analysis, TNF minimally induces HS expression on the surface of WT control macrophages (Fig. 1C). In contrast, RBP-J deficiency enables TNF to markedly increase HS expression on the cell surface (Fig. 1C), paralleling the enhanced osteoclastogenesis observed in *Rbpj^ΔM^* cells (Fig. 1E, and (*27*)). Mass spectrometry quantification of HS levels further supports this observation (Fig. 1D). These findings suggest that RBP-J restrains HS expression in macrophages in response to TNF. To determine whether cell surface HS contributes to the enhanced osteoclastogenesis caused by RBP-J deficiency, we treated cultures with Heparin Lyase III (HLIII), which specifically digests HS, or with Protamine, which blocks HS binding to HS-binding proteins. Removal of HS or inhibition of HS-binding activity abolished the elevated osteoclast differentiation in *Rbpj^ΔM^* cultures, reducing it to levels comparable to WT controls that were not treated with these inhibitors (Fig. 1E). In contrast, removal of HS significantly elevated TNF-induced expression of the ISGs, such as *Mx1, Ifit1, Mx2, Ifit2, Irf7* and *Cxcl9* in the RBP-J deficient cells (Fig. 1F). Together, these results establish a critical role for HS in RBP-J mediated TNF-induced inflammatory response and osteoclastogenesis.

### HS 2-O-sulfotransferase Hs2st1 is a key effector of RBP-J in TNF-mediated osteoclastogenic and inflammatory responses

Since HS biosynthesis is regulated by HS sulfotransferases, we next examined whether any of these enzymes are directly controlled by RBP-J. In the enriched HS biosynthetic pathways identified from RNAseq data, several HS sulfotransferase genes were upregulated in RBP-J deficient macrophages (Fig. 2A). To determine whether these genes are direct transcriptional targets of RBP-J, we performed ChIP assays. Among the HS biosynthetic enzymes, RBP-J was found to only bind to the transcriptional start site regions of both *Hs2st1* and *Ext2*, with more than fourfold greater binding at the *Hs2st1* locus (Fig. 2B). Given that HS sulfation levels and patterns critically influence HS interactions with binding partners and its biological functions, we next analyzed the composition of sulfated HS disaccharides using mass spectrometry. We found that while the amount of 2-O-sulfated disaccharides (combined mass of △UA2S-GlcNS6S, △UA2S-GlcNS, △UA2S-GlcNAc6S and △UA2S-GlcNAc from Supplementary Table 1) produced by unstimulated WT and RBP-J deficient BMMs are similar, after TNF stimulation, RBP-J deficient BMMs produce 45% more 2-O-sulfated disaccharides than WT BMMs (Fig. 2C, left panel). Similarly, after TNF stimulation, RBP-J deficient BMMs produce 50% more N-sulfated disaccharides than WT BMMs (Fig. 2C, middle panel). In contrast, we did not find a significant difference between 6-O-sulfated disaccharides between TNF-stimulated WT and RBP-J deficient BMMs (Fig. 2C, right panel), which is consistent with the less pronounced upregulation of *Hs6st1* and *Hs6st2* compared to *Hs2st1* and *Ndst1*. Combined, the result of our HS structural analysis is in full agreement with our RNAseq and ChIP data, together suggesting that *Hs2st1* is a major direct target suppressed by RBP-J. As RBP-J deficiency enabled TNF to elevate Hs2st1 expression (Fig. 2D), we next examined whether Hs2st1 functions as a downstream effector of RBP-J. Knockdown of Hs2st1 (Fig. 2E) markedly suppressed TNF-induced osteoclast differentiation (Fig. 2F) and osteoclast marker gene expression (Fig. 2G) in RBP-J deficient cell cultures. On the contrary, Hs2st1 deletion increased the expression of ISGs, such as *Mx1, Ifit1* and *Cxcl10* (Fig. 2H), and inflammatory genes (Fig. 2I), such as *Il1*, *Il6* and *Tnf*, in RBP-J deficient cells in response to TNF. These findings indicate that Hs2st1 is a key functional target of RBP-J in coordinating TNF-mediated inflammatory gene expression and osteoclastogenesis.

**Figure 2.**
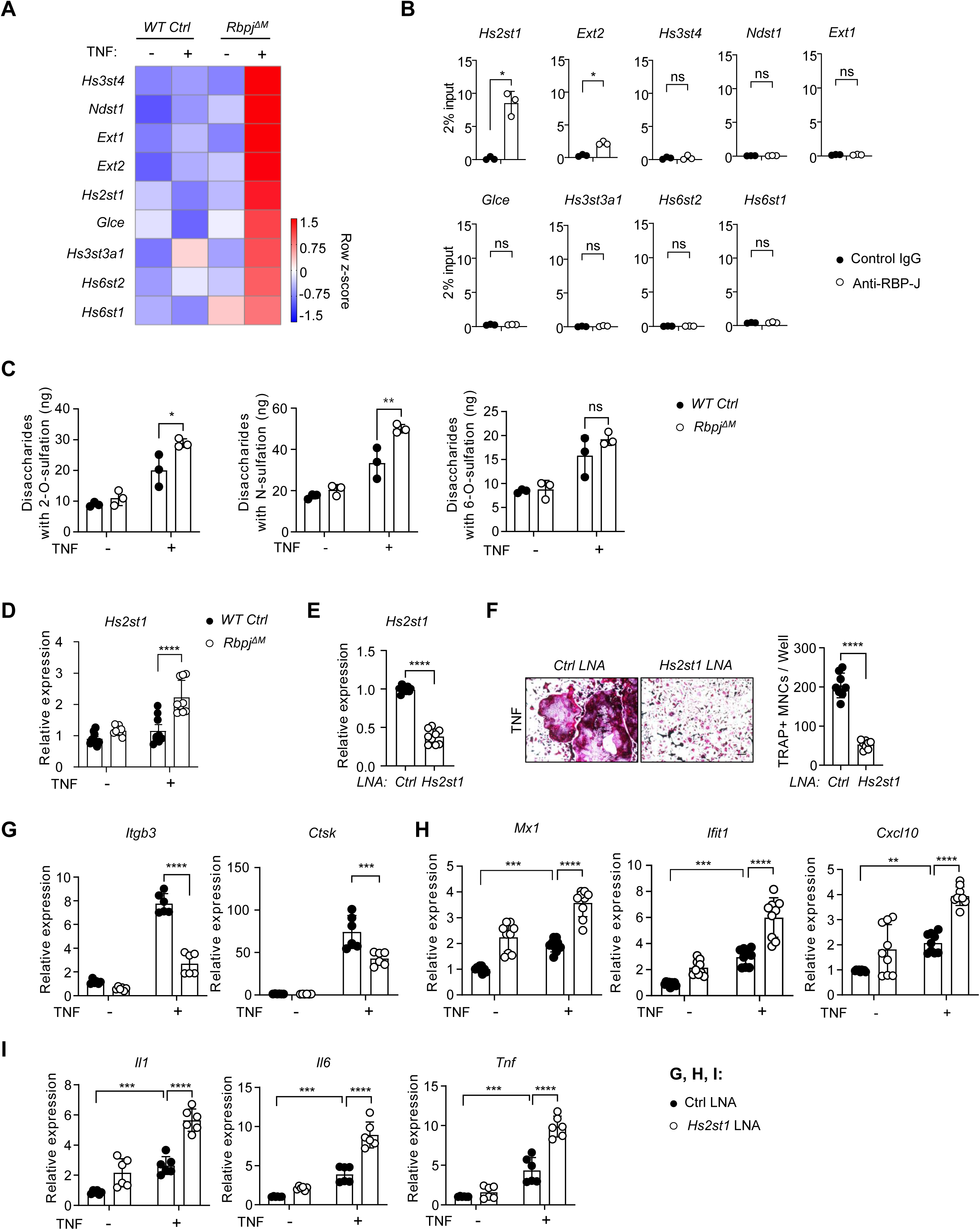
*Hs2st1, a* HS sulfotransferase, is a key RBP-J target that regulates RBP-J activity. (A) Heatmap of gene expression of HS sulfotransferases in wild-type control (WT Ctrl) and *Rbpj^ΔM^* macrophages with or without TNF stimulation for 3 days. (B) ChIP analysis of RBP-J occupancy at the promoter regions of the indicated gene loci of HS sulfotransferases. Control: IgG. n=3. (C) Comparison of total 2-O-sulfated, N-sulfated, and 6-O-sulfated heparan sulfate (HS) disaccharide levels in BMMs from WT Ctrl and *Rbpj^ΔM^* mice treated with or without TNF for 2 days, as measured by LC–MS/MS (n = 3 per group). (D) qPCR analysis of *Hs2st1* expression in the BMMs from WT Ctrl and *Rbpj^ΔM^* mice stimulated with TNF for 2 days. n = 9/group. (E) qPCR analysis of the knockdown efficiency of *Hs2st1* in *Rbpj^ΔM^*BMMs transfected with the *Hs2st1* LNA (40uM) or non-targeting control LNA (Ctrl, 40uM) for 24 hours. (F) Osteoclast differentiation indicated by TRAP staining (left) and the number of TRAP-positive multinucleated cells per well (right) in *Rbpj^ΔM^* BMMs transfected with the *Hs2st1* LNA (40uM), followed by TNF stimulation for 3 days. n = 9/group. (G-I) qPCR analysis of the indicated gene expression in *Rbpj^ΔM^*BMMs transfected with the *Hs2st1* LNA (40uM), followed by TNF stimulation for 1 day (H, n = 9/group), 2 days (I, n = 6/group), or 3 days (G, n = 6/group). Scale bar: F, 100 μm. Data are presented as mean ± SD. *p < 0.05; **p < 0.01; ***p < 0.001; ****p < 0.0001; ns, not statistically significant. Source data are provided as a Source Data file.

### Elevated Hs2st1 enables TNF to effectively drive osteoclast differentiation from macrophages

To further assess the functional role of Hs2st1, we overexpressed Hs2st1 in BMMs using the pMX-puro retroviral transduction system (Fig. 3A, B). Elevated Hs2st1 expression markedly enhanced the capacity of TNF to induce the formation of giant multinucleated osteoclasts (Fig. 3C). Consistently, the expression of the key osteoclastogenic transcription factor *Nfatc1* (encoding NFATc1), along with osteoclast marker genes including *Itgb3* (encoding β3 integrin), *Dcstamp* (encoding DC-STAMP), *Ctsk* (encoding cathepsin K), and *Acp5* (encoding TRAP), was robustly upregulated in Hs2st1 overexpressing macrophages (Fig. 3D). In contrast, TNF-induced ISGs, such as *Mx1*, *Ifit1*, *Cxcl10*, and *Irf7*, were significantly suppressed by Hs2st1 overexpression (Fig. 3E). These findings indicate that elevated Hs2st1 shifts TNF action from inflammatory programs toward osteoclast differentiation in macrophages.

**Figure. 3.**
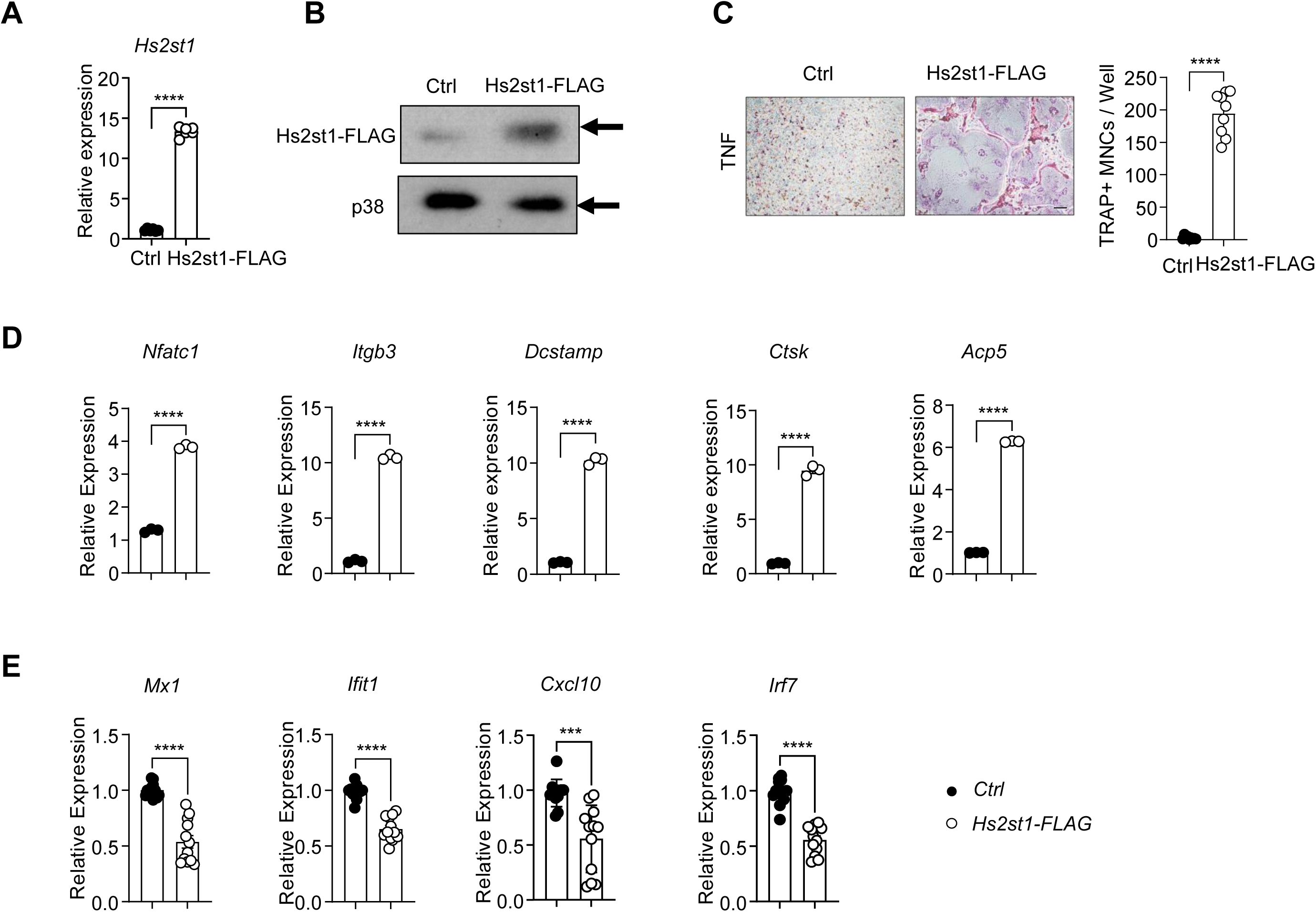
Overexpression of *Hs2st1* promotes TNF-induced osteoclast differentiation of macrophages but decreases ISG expression. (A) qPCR analysis of *Hs2st1* expression (n = 6/group) and (B) Immunoblot analysis of Hs2st1-FLAG expression after BMMs were transduced with retroviral particles encoding *Hs2st1* (pMX-Hs2st1-FLAG, referred to as Hs2st1-FLAG) or the control retroviral particles (pMX empty vector, referred to as Ctrl) for 1 day. Arrows indicate the specific target bands. (C) TRAP staining (Left) and the number of TRAP-positive multinucleated cells per well (Right) and (D) qPCR analysis of osteoclastic gene expression after retroviral transduction followed by TNF stimulation for 3 days (n = 3/group). (E) qPCR analysis of ISG expression after retroviral transduction followed by TNF stimulation for 1 day. (n=12/group), *p<0.05; **p < 0.01; ***p < 0.001; ****p < 0.0001; ns: not statistically significant. Data representative of at least three independent experiments. Data are mean ± SD. Scale bars: C, 100 μm; Source data are provided as a Source Data file.

### Hs2st1 deficiency alleviates inflammatory osteoclastogenesis and arthritic bone erosion

To address the importance of Hs2st1 function in vivo, we generated *Hs2st1* conditional KO mice, in which *Hs2st1* is specifically deleted in myeloid lineage specific macrophages/osteoclast progenitors by crossing *Hs2st1^flox/flox^* mice with *LysMcre* mice (*Hs2st1^f/f^;LysMCre;* hereafter referred to as *Hs2st1^ΔM^*). Sex-matched *LysMcre^+^* littermates served as WT controls (the control). We found that Hs2st1 deficiency in *Hs2st1^ΔM^* BMMs did not impair osteoclast differentiation induced by RANKL (Supplementary Fig.1A). Consistent with this *in vitro* observation, *Hs2st1^ΔM^*mice exhibited no significant changes in bone mass under basal physiological conditions (Supplementary Fig. 1C), and osteoclast formation was comparable between the control and *Hs2st1^ΔM^* mice (Supplementary Fig. 1B). These data indicate that Hs2st1 does not affect physiological RANKL-induced osteoclastogenesis or basal bone remodeling.

We next examined the role of Hs2st1 in TNF-driven responses in vivo using a well-established inflammatory calvarial osteolysis model induced by TNF. Hs2st1 deficiency in *Hs2st1^ΔM^* mice significantly alleviated TNF-induced bone erosion, as shown by μCT analysis of resorption pits on the calvarial bone surface (Fig. 4A), and markedly reduced osteoclast formation in histological sections of calvarial bones compared to WT control mice (Fig. 4B). These findings identify Hs2st1 as a regulator specifically involved in inflammatory bone resorption, with minimal impact on physiological osteoclastogenesis and bone mass.

**Figure 4.**
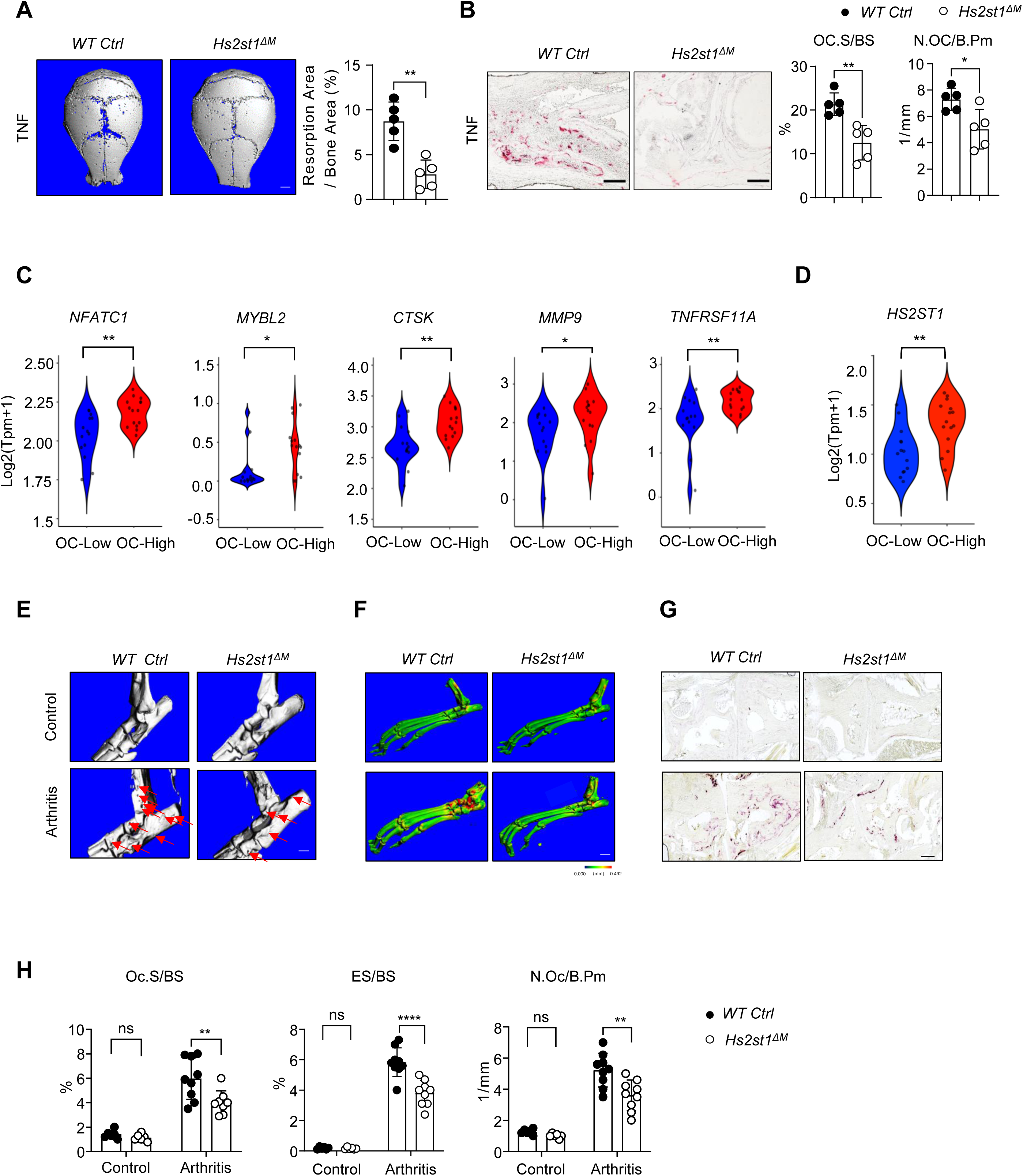
Loss of Hs2st1 attenuates inflammatory osteoclast formation and arthritic bone resorption. (A-B) μCT images (A, Left), the quantification of the resorption area (A, Right), TRAP staining of histological sections (B, Left), and histomorphometric analysis (B, Right) of the slices of the calvarial bones obtained from 12-week-old-male WT and *Hs2st1^ΔM^* mice after PBS or TNF injection to the calvarial periosteum daily for 5 days (n = 5/group). (C) Violin plots showing the expression of the indicated osteoclastic genes in Osteoclastogenic (OC)-High and OC-Low groups. (D) Violin plot showing expression of *HS2ST1* in OC-High versus OC-Low groups. Expression values are shown as log₂(TPM + 1). Each dot represents a single sample. (E) μCT images of the surface of tarsal joints (red arrows: bone erosion). (F) Three-dimensional μCT reconstruction of the hindlimb. TRAP staining (G) and histomorphometric analysis (H) of the tarsal joint sections obtained from the indicated 12-week-old male mice with PBS injection as the control or the littermate male mice that developed K/BxN serum-induced arthritis (Arthritis). n=6/Control group; n=9/Arthritis group. ES/BS, erosion surface/bone surface; Oc.S/BS, osteoclast surface/bone surface; N.Oc/B.Pm, number of osteoclasts per bone perimeter. *p<0.05; **p < 0.01; ***p < 0.001; ****p < 0.0001; ns: not statistically significant. Data are mean ± SD. Scale bars: E, F: 1.0 mm, G: 100 μm. Source data are provided as a Source Data file.

The observation that Hs2st1 promotes inflammatory osteoclastogenesis prompted us to evaluate Hs2st1 expression in rheumatoid arthritis (RA), where TNF is a key cytokine driving both inflammation and bone erosion. We analyzed a recently published dataset (*38*), profiling genome-wide gene expression in synovial monocytes from 29 RA patients. Using an inflammatory osteoclast gene module (*16*), including *NFATC1, MYBL2, CTSK, MMP9,* and *TNFRSF11A*, RA synovial monocyte samples were classified into osteoclastogenic-high (OC-High) and osteoclastogenic-low (OC-Low) groups (Fig. 4C, Supplementary Fig.2). Strikingly, Hs2st1 expression was significantly higher in the OC-High group compared to the OC-Low group (Fig. 4D). Together with our findings that Hs2st1 promotes TNF-induced osteoclastogenesis (Fig. 3), this patient dataset further underscores the disease relevance of Hs2st1 in RA and suggests a positive correlation between Hs2st1 expression and inflammatory osteoclastogenesis and bone erosion.

Guided by this clinical insight, we assessed the role of Hs2st1 in a more pathologically relevant model of inflammatory bone destruction. We employed the K/BxN serum-induced arthritis model (*16, 39, 40*), a well-characterized system for studying inflammatory peri-articular bone erosion mediated by cytokines, such as TNF. Hs2st1 deficiency in *Hs2st1^ΔM^* mice led to striking suppression of peri-articular bone erosion, osteoclast numbers, and resorption surfaces in the tarsal joints (Fig. 4E-H) compared with WT control mice. The clinical course of arthritis, as indicated by joint swelling (Supplementary Fig. 3A, C) and clinical scores (Supplementary Fig. 3B), was not significantly altered in *Hs2st1^ΔM^* mice during the 10-day course, except for the reduced bone erosion. These data suggest that Hs2st1 does not significantly suppress joint inflammation in this model but strongly impacts osteoclast formation and bone destruction.

Taken together, targeted deletion of Hs2st1 alleviates inflammatory osteoclastogenesis and arthritic bone erosion. These findings highlight the therapeutic potential of targeting the Hs2st1 pathway to reduce inflammatory bone loss in diseases, such as RA.

### Loss of Hs2st1 amplifies TNF induced Type-I IFN signaling without altering IFNβ expression

To investigate how Hs2st1 modulates TNF action in macrophages, we performed RNAseq to analyze transcriptomic changes using BMMs from *Hs2st1^ΔM^* and the control mice treated with or without TNF. Reactome pathway and GSEA analyses of the RNAseq data clearly revealed that the type-I IFN signaling pathway was the most significantly enriched TNF-induced pathway in *Hs2st1^ΔM^* cells relatively to the control cells (Fig. 5A, B). The heat map (Fig. 5C) shows that 20 type-I IFN response genes, which are the indicators of the activity of type-I IFN signaling pathway, were remarkably more induced by TNF in the Hs2st1 deficient cells than in the control cells. These findings were further validated using RT-PCR (Fig. 5D). Type-I IFNs includes IFNα and IFNβ. TNF induces the expression of IFNβ but not IFNα in macrophages (*12, 16*). We next examined cellular signaling response to IFNβ, and found that the activation of STAT1/3 was drastically increased by Hs2st1 deficiency (Fig. 5E). Collectively, these results indicate that Hs2st1 deficiency enhances IFNβ signaling. Interestingly, the expression levels of IFNβ induced by TNF are comparable between the *Hs2st1^ΔM^* and the control cells (Fig. 5F). These data indicate that Hs2st1 does not affect the expression of IFNβ but rather its activity/signaling. The IFNβ signaling induced by TNF is a strong autocrine feedback mechanism that regulates cellular functions, such as induction of ISGs and inhibition of osteoclast differentiation (*12, 41*). Since Hs2st1 uniquely controls the 2-O position sulfation of HS, it is highly likely that Hs2st1 suppresses IFNβ activity through HS on the cell surface, thereby limiting IFNβ response and promoting osteoclastogenesis.

**Figure. 5.**
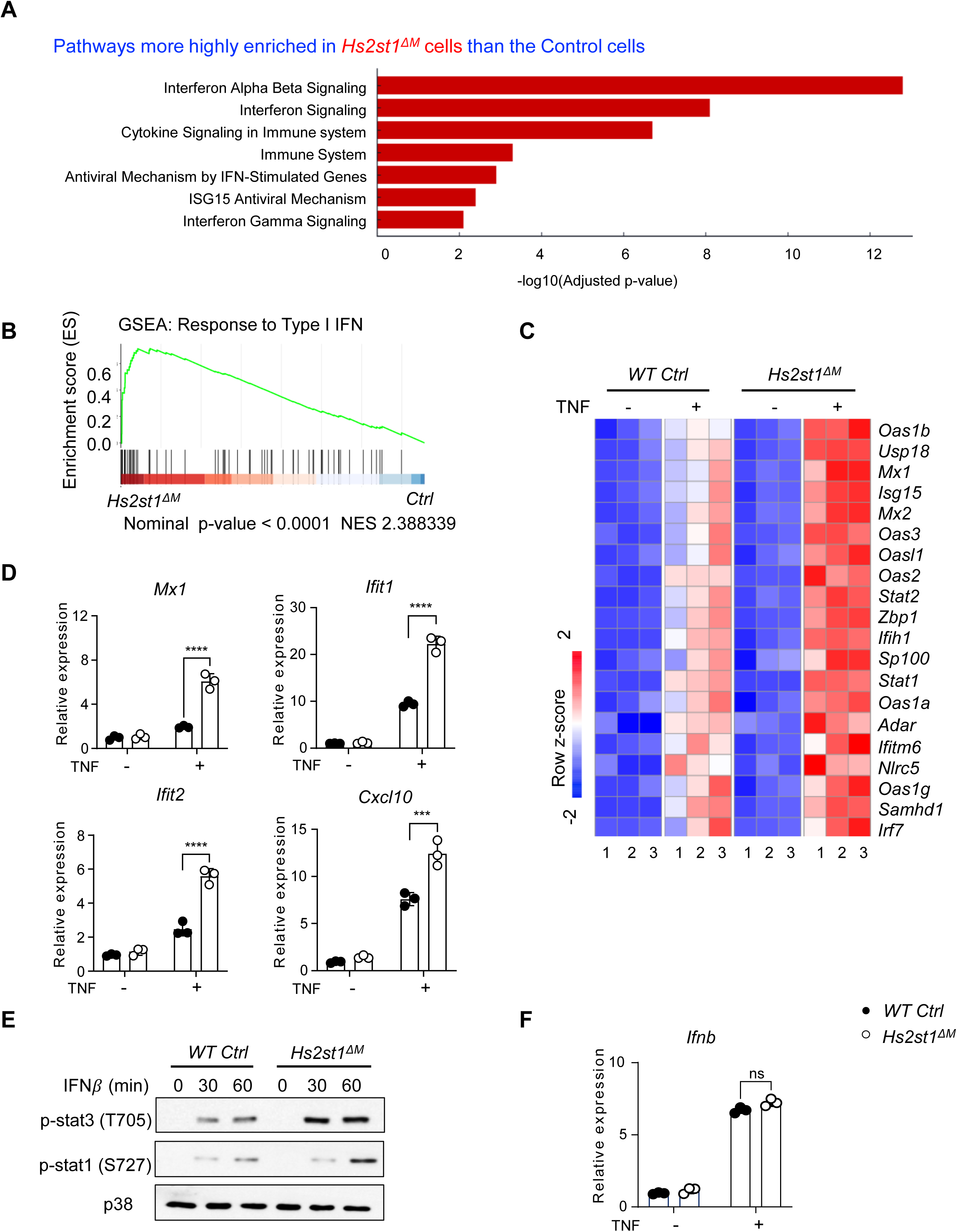
Lack of *Hs2st1* predominantly enhances TNF-induced IFNβ signaling, but does not affect TNF-induced IFNβ expression. (A) Reactome pathway analysis of the 3d-TNF induced DEGs enriched in *Hs2st1^ΔM^* macrophages compared to the WT control cells using EnrichR. Bar length represents –log₁₀(adjusted p value); all shown pathways are significantly enriched in *Hs2st1^ΔM^*cells (*adjusted p < 0.05*). (B) Gene set enrichment analysis (GSEA) of the TNF-induced type I interferon response genes in the control and *Hs2st1^ΔM^* cells. Normalized enrichment score (NES) = 2.39; nominal p < 0.0001. (C) Heatmap of gene expression of the interferon-stimulated genes in the control and *Hs2st1^ΔM^* macrophages with or without TNF stimulation for 3 days. Row z scores of CPMs were shown in the heatmap. n = 3 independent biological replicates for each condition. (D) qPCR analysis of the expression of type-I IFN response genes in BMMs stimulated without or with TNF for 3 days. (E) Immunoblot analysis of phospho-STAT1 (Ser727) and phospho-STAT3 (Tyr705) levels in BMMs treated with IFNβ (100 U/ml) for the indicated times. (F) qPCR analysis of *Ifnb* expression in BMMs after TNF stimulation for 2 days. n=3 / group. **p < 0.01; ***p < 0.001; ****p < 0.0001; ns: not statistically significant. Source data are provided as a Source Data file.

### Hs2st1-dependent 2-O-sulfated HS binds to IFNβ and suppresses IFNβ bioactivity

IFNβ is known to bind HS (*42*), but the binding specificity has never been characterized. Using CHO-K1 cells as a model, we first showed that murine IFNβ binds avidly to CHO-K1 cell surface in a HS-dependent manner (Fig. 6A). By utilizing a widely used Hs2st1-deficient CHO cell line (pgsF), we found that the absence of 2-O-sulfation reduced murine IFNβ binding by 55% (Fig. 6B), suggesting that 2-O-sulfation greatly promotes IFNβ-HS interaction. Of note, we also found that human IFNβ displays a similar binding specificity as murine IFNβ, suggesting a preference for 2-O-sulfated HS is conserved across species (Fig. 6C).

**Figure 6.**
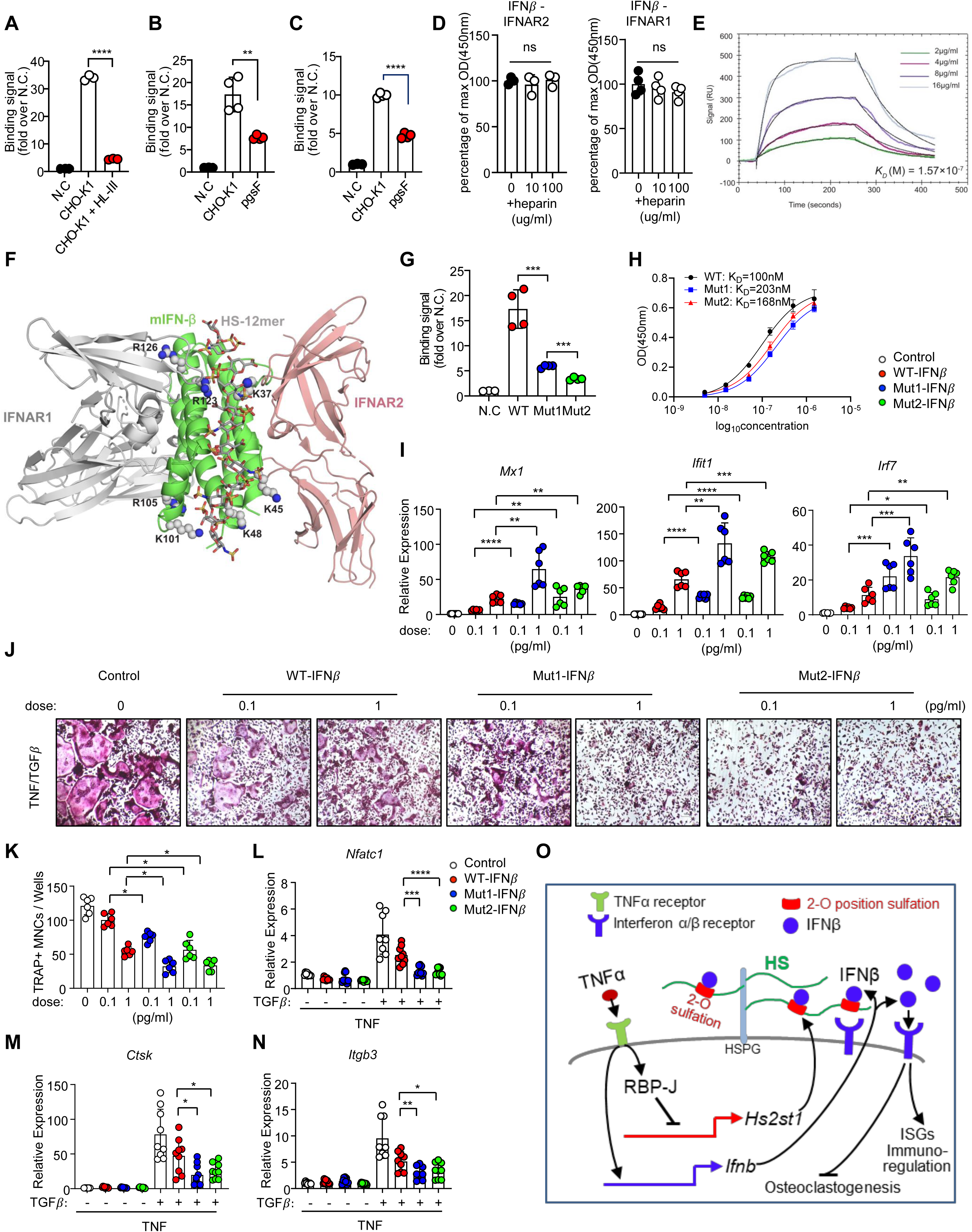
HS binds to IFNβ and finetunes its biological function. (A) Binding of IFN-β to CHO-K1 cells is predominantly mediated by cell surface HS. Murine IFNβ (3 μg/ml) was incubated with untreated CHO-K1 cells or CHO-K1 pretreated with heparin lyase III (HL-III, 5 mU/ml, 15 mins), and the bound IFNβ was detected by a rabbit anti-mIFNβ monoclonal antibody followed by anti-rabbit Alexa-647. Cells stained only with primary and secondary antibodies serve as the negative control (N.C). n = 3. (B) 2-O-sulfated HS promotes binding of murine IFN-β to CHO-K1 cell surface HS. Murine IFNβ (1 μg/ml) was incubated with CHO-K1 cells or pgsF (2-O-sulfation deficient CHO) cells for 1 hr at 4°C, and the bound IFNβ was detected as described above. n = 4. (C) 2-O-sulfated HS promotes binding of human IFN-β to CHO-K1 cell surface HS. Human IFNβ (1 μg/ml) was incubated with CHO-K1 cells or pgsF (2-O-sulfation deficient CHO) cells for 1 hr at 4°C, and the bound IFNβ was detected with a rabbit anti-hIFNβ polyclonal antibody followed by anti-rabbit Alexa-647. n = 3. (D) Heparin does not interfere with IFNβ-IFNAR1 and IFNβ-IFNAR2 interactions. Binding of human IFNβ (200 ng/ml) to immobilized human IFNAR1 and human IFNAR2 in the absence or presence of heparin (10 and 100 μg/ml). Bound IFNβ is detected by a rabbit anti-IFN-β polyclonal antibody. Data representative of two independent experiments. (E) Binding kinetics of mIFNβ-HS interaction. Surface plasmon resonance (SPR) analysis of binding between mIFNβ and immobilized HS dodecasaccharide (GlcNS6S-GlcA-GlcNS6S-(IdoA2S-GlcNS6S)_4_-GlcA). Data representative of three independent experiments. (F) Structural model of IFNβ/IFNAR1/IFNAR2 complex. This model is based on the crystal structures of IFN-α2/IFNAR1/IFNAR2 complex (PDB: 3SE3) and murine IFN-β (mIFNβ) (PDB: 3WCY). IFNAR1 (grey), murine IFN-β (green) and IFNAR2 (salmon) are shown in cartoon representation. HS-binding residues of IFNβ as identified by mutagenesis are displayed in spheres. A 12mer HS oligomer, shown in sticks (carbon backbone in grey, sulfur in yellow, oxygen in red and nitrogen in blue), is manually modeled onto the HS-binding sites to show the potential orientation of bound HS chain. (G) Mut1 mIFNβ and Mut2 mIFNβ show greatly reduced binding to cell surface HS. Binding of WT-, Mut1- and Mut 2-mIFNβ (1 μg/ml) to CHO-K1 cells surface was determined by a FACS-based binding assay. The bound mIFNβ were detected by staining with a rabbit anti-mIFNβ antibody, followed by anti-rabbit IgG Alexa-647. The shaded histogram is from cells stained only with primary and secondary antibodies. Data representative of three independent experiments. (H) Mut1 mIFNβ and Mut2 mIFNβ display similar binding affinity as WT mIFNβ to IFNAR1. Binding of WT-, Mut1- and Mut 2-mIFNβ to immobilized murine IFNAR1 was determined by ELISA-based binding assay. Data representative of two independent experiments. (I) qPCR analysis of the indicated ISG expression in the presence or absence of WT mIFNβ, Mut1 or Mut2 variants at the indicated concentrations for 4 hours (n = 6/group). (J, K) Osteoclast differentiation indicated by TRAP staining (J) and quantification (K) of TRAP-positive multinucleated cells per well (n = 6/group). Scale bar, 100 μm. WT BMMs were treated with TGFβ priming followed by TNF stimulation without or with WT mIFNβ, Mut1 or Mut2 variants at the indicated concentrations for 4 days. (L-N) qPCR analysis of the indicated genes in TGFβ primed BMMs treated without or with TNF (40ng/ml) in the presence or absence of 1 pg/ml of WT mIFNβ, Mut1 mIFNβ or Mut2 mIFNβ for one day (n = 9/group). (O) Schematic illustration showing the RBP-J–Hs2st1–2-O-sulfated heparan sulfate (HS)– IFNβ dependent molecular network that serves as a gatekeeper, balancing macrophage immunoregulatory function and osteoclastogenic potential. *\*p<0.05; **p < 0.01; ***p < 0.001; ****p < 0.0001;* ns, not statistically significant. Data are mean ± SD. Source data are provided as a Source Data file.

To understand the mechanism by which HS inhibits IFNβ signaling through direct binding to IFNβ, we first examined whether HS directly inhibits the interaction between IFNβ and its receptors. We found that binding of IFNβ to immobilized IFNAR2 or IFNAR1 (Fig. 6D) were not affected by heparin (a highly sulfated form of HS), which suggests that binding of IFNβ to HS does not directly interfere with the formation of IFNAR1-IFNβ-IFNAR2 signaling complex. Another potential mechanism by which HS inhibits IFNβ signaling could be that it regulates the bioavailability of IFNβ to its receptors. By surface plasmon resonance (SPR), we first determined the binding kinetics and affinity of IFNβ–HS interaction. Binding of IFNβ to immobilized HS oligosaccharide displayed a rapid on rate (K_a_ (1/M*s) = 7 x10^4^), and a relatively fast off rate (K_d_ (1/s) =1 x 10^-2^), resulting in a binding *K*_D_ of 157 nM (Fig. 6E). This binding characteristic suggests that on cells surface, IFNβ might bounce on and off the highly abundant HS repeatedly before it could encounter IFN receptors, which are usually scarcely expressed. If this is true, then when HS expression is upregulated, the bioavailability of IFNβ to IFN receptors would decrease. To examine whether this is true, we decided to create HS-binding deficient IFNβ mutants and test their signaling potency on BMMs. Due to reduced binding to HS, we predict that these mutants would have improved bioavailability and enhanced IFNβ signaling compared to WT IFNβ.

To this end, we first mapped the HS-binding site of IFNβ by using site-directed mutagenesis. Based on the co-crystal the structure of IFNα2/IFNAR1/IFNAR2 ternary complex (*43*), we first identified 9 basic residues located at the same surface of IFNβ as potential HS-binding residues (Fig. 6F). Alanine mutants of these residues were generated and their binding to heparin sepharose column were examined. We found 7 out of 9 mutants displayed greatly reduced binding to heparin column (Supplementary Table 2), suggesting these residues together form the HS-binding sites of IFNβ. We further prepared several double mutants and found they all resulted in even greater reduction in heparin binding compared to parental single mutants. Among these mutants, we picked K48A (hereafter referred to as Mut1-IFNβ) and K37A-K48A (hereafter referred to as Mut2-IFNβ) for further studies because they displayed the most severe reduction in HS-binding among single and double mutants, respectively (Supplementary Table 2). As expected, binding of Mut1-IFNβ and Mut2-IFNβ to cell surface HS was also greatly impaired, displaying 66% and 81% reduction in binding compared to WT IFNβ, respectively (Fig. 6G). We further determined that Mut1-IFNβ and Mut2-IFNβ displayed similar binding affinity to immobilized IFNAR1 as WT IFNβ (Fig. 6H), which is expected because K37 and K48 are located on a surface not involved in binding to IFNAR1 (Fig. 6F).

IFNβ induces ISGs to regulate macrophage inflammatory and immune responses but also inhibits macrophage differentiation into osteoclasts (*12, 41*). To assess the biological function of IFNβ mutants defective in HS binding, we treated BMMs with these mutants and found that both mutants induced substantially higher expression levels of ISGs, such as *Mx1, Ifit1*, and *Irf7*, compared to WT IFNβ at the same concentrations (Fig. 6I). We then evaluated the ability of these mutants to inhibit osteoclastogenesis using the established TGFβ/TNF-induced osteoclastogenic system in BMMs (*16*). Compared to WT IFNβ, the mutants exhibited much stronger inhibition of osteoclast formation (Fig. 6J, K) and of osteoclastogenic gene expression, including *Nfatc1*, *Itgb3*, and *Ctsk* (Fig. 6L-N). These point-mutation experiments highlight the biological importance of HS in modulating IFNβ activity on macrophage inflammatory responses and osteoclastogenesis. Collectively, these findings establish 2-O-sulfated HS as a biological gatekeeper of IFNβ activity on the cell surface, fine-tuning the abundance of IFNβ-binding sites in balancing macrophage immunoregulatory function and osteoclastogenic potential (Fig. 6O).

## DISCUSSION

Unlike macrophages in other tissues, macrophages in the bone have a unique bipotential. They not only carry out their primary roles in inflammatory response and immune regulation, but also have the ability to differentiate into bone-resorbing osteoclasts. While extensive research has focused on mechanisms that regulate either inflammatory activities or osteoclast differentiation, much less is known about how the bipotential of macrophages is modulated. Certain signaling pathways, such as NF-κB and MAPK, are shared and can promote both functions (*1, 44*). However, little is known about the molecular mechanisms that drive macrophages toward one fate over the other, especially in response to the same stimulus, such as TNF. In inflammatory bone diseases, such as rheumatoid arthritis (RA), where both inflammation and bone resorption are prominent features, understanding these regulatory mechanisms is critical. It would allow for the development of more refined therapeutic strategies that can simultaneously and effectively control both pathological processes.

In this study, we discovered a previously unrecognized pathway mediated by RBP-J-Hs2st1-2-O-sulfated heparan sulfate (HS)-IFNβ axis in response to TNF, which acts as a gatekeeper network to balance macrophage inflammatory activity and osteoclast differentiation. In this network, RBP-J plays a key role in promoting inflammatory responses while inhibiting osteoclastogenesis. We found that Hs2st1 is a novel downstream target suppressed by RBP-J. Hs2st1 inhibits TNF-induced interferon-stimulated gene (ISG) expression while promoting osteoclast differentiation. The RBP-J–Hs2st1 axis triggers a specific inside-out signaling cascade, Hs2st1 → 2-O sulfated-HS → IFNβ, where reduced Hs2st1 expression leads to diminished 2-O-sulfation of HS. This, in turn, facilitates the ‘release’ of IFNβ to activate its receptors, enhancing IFNβ signaling and thereby suppressing osteoclastogenesis. Our results therefore establish 2-O-sulfated HS as a key cell surface gatekeeper that fine-tunes IFNβ activity, modulating macrophage responses and osteoclastogenic potential. When HS binding to IFNβ is weak, such as in healthy individuals or early-stage RA where Hs2st1 expression (Supplementary Fig. 4) and 2-O-sulfation are low (*45*), macrophages are more likely to express ISGs and suppress osteoclastogenesis in response to TNF. In contrast, when HS binding is strong, such as in later stages of RA where Hs2st1 expression (Supplementary Fig. 4) and 2-O-sulfation are elevated (*45*), macrophages are more prone to differentiate into osteoclasts with reduced ISG expression. These findings reveal a fine-tuning mechanism with important implications for developing more precise and effective treatments for inflammatory diseases with bone resorption.

How HS interacts with IFNβ and by what mechanisms HS regulates IFNβ bioactivity/signaling was poorly understood. Although a prior study showed that HS-IFNβ interaction interfered IFNβ signaling, it did not reveal the molecular mechanisms by which HS interferes IFNβ signaling, nor did it determine the HS-binding sites of IFNβ (*42*). Our findings revealed that HS does not directly interfere with the binding of IFNβ to its receptors. This indicates that the inhibitory effect of HS is not mediated by direct competition with IFNAR1 or IFNAR2 for IFNβ binding, and that the HS-binding sites on IFNβ are spatially separated from the molecular surfaces required for interaction with IFNAR1 and IFNAR2. With this structural insight, we successfully identified several key residues on IFNβ responsible for binding to HS and prepared two HS-binding deficient IFNβ mutants. With these mutants, we provided further confirmation of the critical role of IFNβ-HS interaction in regulating macrophage immune regulation and cell fate of osteoclast differentiation. We believe IFNβ-HS interaction holds potential as a novel therapeutic target for modulating macrophage function in disease contexts, particularly inflammatory diseases with bone defects. Type I IFNs, including both IFNβ, IFNκ and IFNα family, share structural homology and signal through the same receptor complex, although with different binding affinities. To the best of our knowledge, among type I IFNs, IFNκ has also been shown to bind heparin (*46*), while IFNα4 shown no binding to HS (*42*). This difference in HS-binding capacity is likely attributable to their relatively low sequence homology (approximately 31–38%, (*47*)) between IFNβ and IFNα. However, whether other members of IFNα family interact with HS remains unknown (*48*). Beyond type I IFNs, the type II IFN, IFNγ, has also been shown to interact with HS. Interestingly, the binding of HS appears to enhance IFNγ activity (*49*), which contrasts with our mechanistic model of IFNβ, where HS binding suppresses its activity. While type I and type II IFNs play distinct dominant roles across various biological contexts and diseases, both are key regulators of immune responses and inflammation. The modulation of their activity by HS may offer a strategy to selectively target these cytokines for therapeutic purposes. For instance, in diseases like multiple sclerosis, where exogenous IFNβ is used therapeutically to suppress autoimmunity, strategies that enhance IFNβ signaling, such as limiting its inhibition by HS or using IFNβ mutants with reduced HS binding generated in this study, may improve its clinical efficacy. Similar strategies may be applied to antiviral infections by enhancing IFNβ activity through its interaction with HS.

Sulfation at the 2-O position of HS can only be achieved through the enzymatic activity of Hs2st1, which has only a single isoform in all animal species sequenced so far. This is in sharp contrast to all other families of HS sulfotransferases (Ndst, Hs6st and Hs3st), where multiple isoforms are found to perform the same enzymatic reaction (*20*). As a result, alteration of the expression level of Hs2st1 often incurs a direct and predictable structural changes of HS, which can result in dramatic changes in its binding to specific HS-binding proteins. For instance, binding of osteoprotegerin (OPG) to HS critically depends on 2-O-sulfated HS, and Hs2st1-deficient osteoblasts have greatly impaired capacity to retain OPG on its surface to efficiently inhibit RANKL (*26, 50*). Compared to OPG, the binding specificity of IFNβ to HS is less stringent, as Hs2st1-deficient CHO cells still retain 40-50% of binding capacity to IFNβ. However, even this partial specificity for 2-O-sulfation is sufficient to impact the bioactivity of IFNβ. The sensitivity of this HS-binding dependent regulation of IFNβ is even more remarkable considering that a mere 45% increase in 2-O-sulfated HS is sufficient to switch the fate of TNF treated BMMs from inflammatory state to osteoclastogenesis. Our finding provides a prime example of how moderate changes of HS structure can have profound biological consequences.

The effects of type I IFNs on inflammatory regulation in RA have been mixed (*51, 52*), with some studies reporting that type I IFNs promote inflammation, while others show the opposite. In our RA mouse model, Hs2st1 deficiency did not significantly alter the inflammatory response. Since Hs2st1 regulates 2-O-sulfation and thereby modulates IFNβ activity, our results suggest that HS-mediated IFNβ activity may not have a major impact on inflammation in RA. One possible explanation is a threshold effect, where the changes in IFNβ activity caused by HS modulation are not sufficient to influence the overall inflammatory state. Another likely explanation is that RA associated inflammation is shaped by a complex network of factors, including contributions from multiple cell types, cytokines, and chemokines. This intricate inflammatory environment could override the effects of HS-mediated IFNβ immunoregulatory activity observed in macrophages. In contrast, the impact of the Hs2st1-2-O-HS-IFNβ axis on osteoclastic bone erosion in RA was striking. Hs2st1 deficiency strongly suppressed osteoclast formation and bone resorption in the RA model, indicating that HS-mediated IFNβ activity is more sensitive in regulating osteoclastic suppression than in modulating inflammation. Over the course of RA progression, particularly in the late stages, osteoclast-driven bone erosion becomes a major cause of bone destruction, joint deformity, and loss of function. Therefore, targeting the Hs2st1-2-O-HS-IFNβ axis may represent a promising complementary therapeutic approach that can inhibit inflammatory osteoclastogenesis and bone erosion without compromising the overall immune response.

The RBP-J-HS network appears not to affect physiological osteoclastogenesis or bone remodeling, as both RBP-J deficient mice (*14, 27*) and Hs2st1-deficient mice show no bone defects. Consistent with this phenotype, basal osteoclastogenesis driven by RANKL signaling remains normal in these mice. Because maintaining basal osteoclastogenesis and bone remodeling is essential to avoid adverse effects from treatments (*17, 18*), targeting this network could provide a novel therapeutic strategy that selectively inhibits inflammatory bone resorption while maintaining physiological bone homeostasis. Interestingly, we found that both WT and mutant IFNβ inhibited RANKL-induced osteoclastogenesis to a similar extent in vitro (Supplementary Fig. 5). This observation aligns with and may explain the normal basal osteoclast formation and bone mass in both RBP-J and Hs2st1 deficient mice. This is presumably because RANKL is a much stronger osteoclastogenic stimulus than TNF, often requiring higher levels of IFNβ to counteract its effects. The difference in the inhibition of inflammatory osteoclastogenesis between WT and mutant IFNβ was also dose-dependent. For example, at low concentrations (0.1-1 pg/ml), the mutant showed elevated osteoclastic inhibitory activity compared with WT, but at ≥10 pg/ml, the difference disappeared. This effect is expected because when the amount of IFNβ in the system is sufficiently abundant, the chance of some of them reaching the receptors would greatly increase. Taken together, these IFNβ mutation experiments establish a proof-of-concept model for investigating the newly identified RBP-J-HS mediated network regulating inflammation and osteoclastogenesis in macrophages. Future work will focus on designing monoclonal antibodies or HS oligosaccharides that target the HS-binding sites of IFNβ. These studies will not only advance our biochemical understanding but also lay the groundwork for translational strategies aimed at modulating IFNβ activity in inflammatory diseases.

## METHODS

### Animal study and inflammatory bone resorption models

We generated mice with myeloid/macrophage-specific deletion of *Hs2st1* by crossing the *Hs2st1* ^flox/flox^ mice (gift from Jeffery Esko, University of California San Diego, La Jolla, CA) with the mice carrying a lysozyme M promoter-driven Cre transgene on the C57BL/6 background (LysMcre; The Jackson Laboratory, Stock No: 004781). Sex- and age-matched *Hs2st1* ^flox/flox^; LysMcre(+) mice (referred to as *Hs2st1^ΔM^*) mice and their littermates with *Hs2st1*^+/+^; LysMcre(+) genotype as WT controls (hereafter referred to as WT Ctrl) were used for experiments. Similarly, mice with myeloid/macrophage-specific deletion of *Rbpj* were generated by crossing the *Rbpj*^flox/flox^ mice with the LysMcre mice. Sex- and age-matched *Rbpj*^flox/flox^; LysMcre(+) mice (referred to as *Rbpj^ΔM^*) and their littermates with *Rbpj*^+/+^; LysMcre(+) genotype as WT controls (hereafter referred to as WT Ctrl) were used for experiments. All mice were housed under a 12-hour light/dark cycle at room temperature, with ad libitum access to dry laboratory chow and water. All animal procedures were approved by the Institutional Animal Care and Use Committees (IACUC) of the Hospital for Special Surgery and Weill Cornell Medical College.

For inflammatory osteolysis experiments, we employed an established TNF-induced supracalvarial osteolysis mouse model with minor modifications (*16, 41, 53*). Briefly, TNF was administered daily at a dose of 75 μg/kg to the calvarial periosteum of age- and sex-matched mice for 5 consecutive days. Mice were then sacrificed, and calvarial bones were collected for micro-computed tomography (μCT) analysis, sectioning, tartrate-resistant acid phosphatase (TRAP) staining, and histological evaluation.

For inflammatory arthritis experiments, we utilized the K/BxN serum transfer-induced arthritis model (*16, 39*). Pooled K/BxN serum was prepared, and arthritis was induced by intraperitoneal injection of 150 μl serum into male mice on days 0 and 2. Arthritis progression was monitored by measuring the thickness of both wrist and ankle joints using a digital slide caliper (Bel-Art Products). Joint thickness for each animal was calculated as the sum of both wrists and ankles, and joint thickness was represented as the average for each group. Semiquantitative clinical scores were assigned for each of the four limbs according to the following criteria: 0 = no observable swelling; 1 = involvement of a single digit or mild swelling of the foot and ankle with preservation of the normal V-shaped contour; 2 = parallel alignment of the long edges of the foot with loss of the normal V-shape; and 3 = inversion of the V-shape due to expansion of the ankle and hindfoot exceeding the width of the forefoot. Scores from all four limbs were summed to obtain the clinical score (maximum score = 12) as described previously (*54*). Mice were sacrificed on day 10, and serum and paw tissues were collected. Hind paws were processed for μCT analysis, sectioning, TRAP staining, and histological assessment. μCT scans were performed using a Scanco μCT-35 scanner (SCANCO Medical) to evaluate bone volume and 3D bone architecture in accordance with the manufacturer’s instructions and the American Society for Bone and Mineral Research (ASBMR) guidelines (*55*).

### Reagents

Murine M-CSF, murine TNFα, Murine RANKL were purchased from PeproTech. Murine TGFβ1 was purchased from R&D systems. Protamine Sulfate salt from salmon (P4020-1G) was purchased from Sigma-Aldrich.

### Cell culture

Mouse bone marrow cells were harvested from the tibiae and femora of age- and sex-matched mutant and control mice and cultured for 3 days in α-MEM medium supplemented with 10% fetal bovine serum (FBS), glutamine (2.4 mM; Thermo Fisher Scientific), penicillin–streptomycin (Thermo Fisher Scientific), and recombinant murine M-CSF (20 ng/ml). The attached bone marrow–derived macrophages (BMMs) were scraped, seeded at a density of 4.5 × 10^4^ cells/cm², and cultured overnight in α-MEM containing 10% FBS, 1% glutamine, and conditioned medium. Cells were then treated with or without TNFα (40 ng/ml) or RANKL (40ng/ml) in the presence of recombinant mouse M-CSF for the durations indicated in the figure legends. Culture media were exchanged every 3 days. For TGFβ priming/TNF cell cultures, mouse BMMs were cultured for 3 days in α-MEM medium containing 10% FBS, glutamine (2.4 mM, Thermo Fisher Scientific), Penicillin–Streptomycin (Thermo Fisher Scientific) and recombinant murine M-CSF (20 ng/ml) with murine TGFβ1 (1 ng/ml; R&D systems, Minneapolis, MN). The attached BMMs were washed, scraped, seeded at a density of 4.5 × 10^4^/cm^2^, and cultured in α-MEM medium with 10% FBS, 1% glutamine and recombinant murine M-CSF for overnight. The cells were then treated with TNF (40 ng/ml) and recombinant murine M-CSF (20 ng/ml) in the presence or absence of WT, Mut1 mIFNβ and Mut2 mIFNβ indicated in the figure legends. Culture media were exchanged every 3 days. TRAP staining was performed using an acid phosphatase leukocyte diagnostic kit (Sigma-Aldrich) according to the manufacturer’s instructions. TRAP-positive cells were identified by characteristic red staining.

### In vitro gene silencing

Antisense inhibition using locked nucleic acid (LNA) technology (Qiagen) was employed to silence gene expression in vitro. LNA oligonucleotides specifically targeting *Hs2st1* or non-targeting control LNAs (Qiagen) were transfected into mouse bone marrow–derived macrophages at a final concentration of 40 nM using TransIT-TKO transfection reagent (Mirus) according to the manufacturer’s instructions.

### Retroviral gene transduction

Retrovirus packaging was performed by transfecting the retroviral vectors pMX-control (Cell Biolabs) or pMX–murine *Hs2st1* into Plat-E cells (Cell Biolabs) using FuGENE 6 (Promega), as described previously (*56*). Bone marrow–derived macrophages (BMMs) were infected with retroviral supernatants in the presence of 8 μg/ml polybrene for 48 hours. The culture medium was then replaced with fresh medium for at least 5 hours before stimulation with TNFα (40 ng/ml) in the presence of recombinant mouse M-CSF. Culture media were exchanged every 3 days. The cells were harvested on Day 1 or Day 3 after treatment, and total RNA was extracted for quantitative PCR (qPCR) analysis.

### Reverse transcription and real-time PCR

DNA-free RNA was extracted from cells using the RNeasy Mini Kit (Qiagen) with on-column DNase treatment. Reverse transcription was performed using 1 μg of total RNA, random hexamers, and M-MLV Reverse Transcriptase (Thermo Fisher Scientific) according to the manufacturer’s instructions. Quantitative real-time PCR (qRT-PCR) was carried out in triplicate on a QuantStudio 5 Real-Time PCR System using Fast SYBR® Green Master Mix (Thermo Fisher Scientific) and 500 nM primers (*57*). mRNA levels were normalized to Gapdh as the endogenous control. The primer sequences used for qRT-PCR were as follows*: Hs2st1*: 5’-AGAAAGGGCAATTGCAAGGC-3’ and 5’ACGAGGTGCTTGCAGTTTTG-3’*; Nfatc1*: 5′-CCCGTCACATTCTGGTCCAT-3′ and 5′-CAAGTAACCGTGTAGCTCCACAA-3′; *Acp5*: 5′-ACGGCTACTTGCGGTTTC-3′ and 5′-TCCTTGGGAGGCTGGTC-3′; *Ctsk:* 5′-AAGATATTGGTGGCTTTGG-3′ and 5′-ATCGCTGCGTCCCTCT-3′; *Itgb3*: 5′-CCG GGGGACTTAATGAGACCACTT-3′ and 5′-ACGCCCCAAATCCCACCCATACA-3′; *Dcstamp:* 5′-TTTGCCGCTGTGGACTATCTGC-3′ and 5′-AGACGTGGT TTAGGAATGCAGCTC-3′; *Mx1*: 5′-GGCAGACACCACATACAACC-3′ and 5′-CCTCAGGCTAGATGGCAAG-3′; *Mx2:* 5’-ACGAGAATTGCCAGGGTTTG-3’ and 5’-ATTTCAGTGACCGTGTGCAG-3’; *Ifit1*: 5′-CTCCACTTTCAGAGCCTTCG-3′ and 5′-TGCTGAGATGGACTGTGAGG-3′; *Ifit2:* 5′-AAATGTC ATGGGTACTGGAGTT-3′ and 5′-ATGGCAATTATCAAGTTTGTGG-3′; *Irf7:* 5’-CAGCGAGTGCTGTTTGGAGAC-3’ and 5’-AAGTTCGTACACCTTATGCGG-3’*; Cxcl9:* 5’-TTTGGGGTTCTACAGTGGAG-3’ and 5’-GAAGATGGGATCAAGTTAATA-3’*; Cxcl10:* 5’-ATTCTTTAAGGGCTGGTCT-3’ and 5’-CACCTCCACATAGCTTACA-3’*; Ifnb:* 5’-ATGAGTGGTGGTTGCAGGC-3’ and 5’-TGACCTTTCAAATGCAGTAGATTC-3’; *Il1b:* 5’-AGCTTCCTTGTGCAAGTGTCT-3’ and 5’-GACAGCCCAGGTCAAAGGTT-3’; *Il6*: 5’-TACCACTTCACAAGTCGGAGGC-3’ and 5’-CTGCAAGTGCATCATCGTTGTTC-3’; *Tnf:* 5’-CCCTCACACTCAGATCATCTTCT-3’ and 5’-CTTTGAGATCCATGCCGTTG −3’*; Gapdh*: 5′-ATCAAGAAGGTGGTGAAGCA-3′ and 5′-AGACAACCTGGTCCTCAGTGT-3′.

### Immunostaining

Bone marrow–derived macrophages (BMMs) were cultured on chambered coverslips (CultureWell; Invitrogen) and fixed with 4% paraformaldehyde (PFA). Immunofluorescence staining was performed using HS20 (*58*), a human anti-HS mAb (0.1 μg/mL; BE0412, Bio X Cell) overnight, followed by goat anti-human IgG (H+L) cross-adsorbed secondary antibody conjugated to Alexa Fluor™ 594 (Invitrogen). Nuclei were counterstained with DAPI. Fluorescence images were acquired using a Zeiss fluorescence microscope, and channels were merged using Zeiss imaging software.

### Immunoblot analysis

Total cellular extracts were prepared using lysis buffer containing 150 mM Tris-HCl (pH 6.8), 6% SDS, 30% glycerol, and 0.03% bromophenol blue; 10% 2-mercaptoethanol (2-ME) was added immediately before harvesting the cells. Lysates were separated on 7.5% SDS–PAGE gels, transferred to Immobilon-P membranes (Millipore), and probed with specific primary antibodies. Protein bands were visualized using Western Lightning Plus-ECL detection reagent (PerkinElmer). The following primary antibodies were used: anti–p38α (sc-535, 1:3000; Santa Cruz Biotechnology), anti–phospho-STAT1 (Tyr701) (9177, 1:1000; Cell Signaling Technology), and anti–phospho-STAT3 (Tyr705) (9131S, 1:1000; Cell Signaling Technology). The anti-rabbit IgG, HRP-linked secondary antibody (7074, 1:3000; Cell Signaling Technology) was used to detect these primary antibodies in the experiments.

### Chromatin Immunoprecipitation (ChIP) Assay

BMMs (1×10⁷ cells per condition) from WT control mice were crosslinked with 0.8% formaldehyde for 10 minutes at room temperature, followed by quenching with 125 mM glycine for 5 minutes. Nuclei isolation and chromatin digestion were performed using the SimpleChIP® Enzymatic Chromatin IP Kit (Cell Signaling Technology, #9003) according to the manufacturer’s instructions. The digested chromatin was further sonicated using a Bioruptor® Pico (Diagenode, NJ, USA) for six cycles (30 seconds on/30 seconds off). Chromatin digestion efficiency and DNA concentration were assessed by agarose gel electrophoresis and quantified using a NanoDrop spectrophotometer, respectively. Chromatin lysates were incubated overnight at 4°C with an anti–RBP-J antibody (Cell Signaling Technology, #5313; 1:10) or an equivalent amount of normal rabbit IgG (Cell Signaling Technology, #2729) as an isotype control, followed by immunoprecipitation with ChIP-grade Protein G magnetic beads (Cell Signaling Technology, #9006) for 6 hours at 4°C. After cross-link reversal by overnight incubation at 65°C in the presence of 0.2 M NaCl and Proteinase K treatment (Cell Signaling Technology, #9003; 20 mg/ml), chromatin DNA was purified using the QIAquick PCR Purification Kit (Qiagen, #28104). Chromatin DNA was analyzed by qPCR. RBP-J occupancy was calculated relative to input DNA and normalized to the IgG control (*34*). The qPCR primers covering promoter regions used in the ChIP assay were as follows: *Hs2st1*: 5’-TGGCGATGCTCTTCTTGGAG-3’ and 5’-AAGCCCACTTGCTATAGCCG-3’; *Ext2*: 5’-TGACTCCTCCTGTCCAGCAG-3’ and 5’-CCACGGCTGTGTCGGATC-3’; *Hs3st4*: 5’-GAGTCGCTCACCACTGACAA-3’ and 5’-GTGACGAATGAGGCTCCCAA-3’; *Ndst1*:5’-CCTGTTCCTGCTGTTCGTCT-3’ and 5’-CCTGGCCCAGCTGTGAATAA-3’; *Ext1*: 5’-CGGACTGGAGCTGAAAGTGT-3’ and 5’-TTCCGACCACTGTGCTCTTC-3’; *Glce*: 5’-CCAGCGACAAAGCAATCCAG-3’ and 5’-AGCCCTAACACCTTGCTTCC-3’; *Hs3st3a1*: 5’-AACGCAAAGGAACGGGAGAT-3’ and 5’-CGACTTTGGCTGGGATGGAT-3’; *Hs6st2*: 5’-AGGCTCGCTCCTGATCCTTA-3’ and 5’-CGAGAGACACACCCCTAGGA-3’; *Hs6st1*: 5’-GCATGAATGGCCGGATGAAC-3’ and 5’-CTGGAGAGCGCCAAGAAGAA-3’.

### RNAseq and analysis

Total RNA was extracted from the cultured mouse primary bone marrow derived macrophages using the RNeasy Mini Kit (Qiagen) according to the manufacturer’s instructions. Poly(A)+ transcripts were purified, and sequencing libraries with multiplexed barcode adapters were prepared using the NEBNext Ultra II RNA Library Prep Kit for Illumina (NEB) following the manufacturer’s protocol. All samples passed quality control assessment using a Bioanalyzer 2100 (Agilent). High-throughput sequencing (50 bp, paired-end) was performed on an Illumina HiSeq 4000 platform at the Weill Cornell Medicine Genomics Resources Core Facility, achieving a sequencing depth of 30–50 million reads per sample.

RNAseq reads were aligned to the murine genome (GRCm38/mm10) using HISAT2 with default parameters (*59*). Read counts were obtained using HTSeq-count, and transcript abundances (counts per million, CPM) were calculated with edgeR (*60*). Genes with low expression (<1 CPM) across all conditions were excluded from downstream analyses. Differentially expressed genes (DEGs) were identified using edgeR on three independent biological replicates, with Benjamini–Hochberg false discovery rate (FDR) correction applied for multiple testing. Genes with adjusted *p* value < 0.05 and fold-change (FC) > 1 (Fig. 5a) or FC > 2 (Fig. 1A, B) were considered significantly differentially expressed. Pathway enrichment analyses were performed using clusterProfiler and EnrichR, and gene set enrichment analysis (GSEA) was conducted with clusterProfiler. All violin plots, bar plots, and heat maps were generated using the ggplot2 package in R.

### Analysis of gene expression from RA samples

Bulk RNAseq data from sorted human monocytes in Figure 4C, D were obtained from the ImmPort repository (https://www.immport.org/shared/study/SDY998; study accession code SDY998), which includes RNAseq profiles of sorted human monocytes (*38*) and a preprocessed gene expression matrix expressed in transcripts per million (TPM) units. To evaluate osteoclastogenesis-associated gene set enrichment, we transformed the TPM matrix to log₂(TPM + 1) and performed gene set variation analysis (GSVA) using the GSVA R package. A predefined osteoclastogenic (OC) signature gene set comprising *NFATC1, MYBL2, CTSK, MMP9,* and *TNFRSF11A* (*16*) was used. GSVA scores were calculated to reflect the relative enrichment of this OC gene set in each sample. Samples were then stratified into osteoclastogenic-high (OC-High) and osteoclastogenic-low (OC-Low) groups based on their GSVA enrichment scores, using the median GSVA value as the threshold. Samples with scores greater than or equal to the median (≥ median) were classified as OC-High, and those with scores below the median (< median) were classified as OC-Low. The resulting GSVA scores and corresponding gene expression levels were visualized using violin plots generated with the *ggplot2* package (version 3.4.4) in R, displaying both the kernel density distributions and individual sample points. Statistical comparisons of gene expression between groups were performed using pairwise Wilcoxon rank-sum tests, with significance indicated as * *p* < 0.05, ** *p* < 0.01.

RNAseq data from synovial biopsy samples in Supplementary Figure 4 were obtained from the Gene Expression Omnibus (GEO; accession number GSE89408), a bulk RNAseq study of synovial tissue from patients with and without RA (*61*). Transcript abundances (counts per million, CPM) were calculated with edgeR. Normalized log₂ (CPM+1) values were used for visualization of *HS2ST1* expression across clinical groups (Healthy, Early RA, and Established RA) as defined in the original metadata of the study. Violin plots were generated using the ggplot2 package (version 3.4.4) in R, showing both kernel density distributions and individual sample points. Statistical comparisons of gene expression between groups were performed using pairwise Wilcoxon rank-sum tests, with significance indicated as * *p* < 0.05, ** *p* < 0.01, **** *p* < 0.0001.

### Expression and purification of murine IFNβ in 293-F cells

Complete open reading frame of murine IFNβ (Genscript) was cloned into pcDNA3.1 (Thermo Fisher Scientific) expression vector and the sequence of the final plasmid was confirmed by sanger sequencing. Recombinant murine IFNβ was expressed in 293-Freestyle cells (Thermo Fisher Scientific) by transient expression using FectoPRO transfection reagent (Polyplus transfection). Purification of IFNβ from culture supernatant was carried out using HiTrap heparin-Sepharose column at pH 7.1 (HEPES buffer) followed by Superdex 200 Increase size exclusion column using an AKTA Go protein purification system. After purification, IFNβ was >99% pure as judged by silver staining. The endotoxin level of the purified IFNβ was < 0.01EU/µg as determined by ToxinSensor Chromogenic LAL Endotoxin Assay Kit (Genscript).

### Surface plasma resonance (SPR)

SPR was performed on an OpenSPR instrument (Nicoya). Biotinylated 12merNS2S6S (Glycan Therapeutics) was immobilized to a streptavidin sensor chip (Nicoya) based on the manufacturer’s protocol. Briefly, the biotin-12mer conjugate (20 µg/ml) in 150 µl HBS-running buffer (25 mM HEPES, pH 7.1, 0.15M NaCl, 0.05% Tween-20) was injected to channel 2 of the flow cell of the sensor chip at a flow rate of 20 µl/min. The flow cell channel 1 without any immobilization was served as a background control. Different dilutions of IFNβ (concentrations from 2-16 µg/ml) in HBS-running buffer were injected at a flow rate of 20 µl/min. At the end of the sample injection, the sensor surface was washed with HBS to facilitate dissociation. After a 5-min dissociation time, the sensor surface was regenerated by injecting with 150 µl regeneration buffer (0.025M HEPES, pH 7.1, 2M NaCl) at a flow rate of 150 µl/min to get a fully regenerated surface. The sensorgrams were fit with 1:1 Langmuir binding model from TraceDrawer 1.9.2.

### Site-directed mutagenesis

Murine IFNβ mutants were prepared by site-directed mutagenesis and the mutations were confirmed by Sanger sequencing. Expression and purification of the mutants were carried out the same way as the WT IFNβ.

### Heparin–Sepharose chromatography

To characterize the binding of WT IFNβ and IFNβ mutants to heparin, 50 μg of purified IFNβ was applied to a 1 ml HiTrap heparin–Sepharose column (Cytiva) and eluted with a salt gradient from 150 mM to 1 M NaCl at pH 7.1 in 25 mM HEPES buffer. The conductivity measurements at the peak of the elution were converted to the concentration of NaCl based on a standard curve.

### Fluorescence-Activated Cell Sorting

CHO-K1 cells or pgsF cells (2-O-sulfation deficient CHO cell mutant), were lifted from culture dish using EDTA and incubated with WT or mutant IFNβ at 1 µg/mL or 3 µg/ml in PBS with 0.1% BSA for 1 hr at\ 4 °C. Bound IFNβ was stained with monoclonal rabbit anti-mouse IFNβ (1: 1000, Cell Signaling #97450) for 1 hr at 4 °C, followed by goat anti-rabbit IgG-Alexa 647 (1:1,000; ThermoFisher Scientific) for 30 min and analyzed by flow cytometry. For negative control, cells were stained with primary and secondary antibodies only without added IFNβ. In some experiments, cells were pretreated with recombinant HL-III (5 milliunits/ml, produced in our lab) for 15 min at room temperature prior to binding experiments.

### Binding of IFNβ to immobilized IFN receptors

For determination of the effect of heparin on IFNβ/IFNAR1 and IFNβ/IFNAR2 interactions, 100 ng of human IFNAR1-Fc (Acro Biosystems, IF1-H5253) or IFNAR2-Fc (Acro Biosystems, IF2-H5255) were immobilized on to 96 well plate. For IFNβ/IFNAR1 interaction, binding of IFNβ was tested at 10 ng/ml in the presence or absence of 10 or 100 µg/ml heparin. For IFNβ/IFNAR2 interaction, binding of IFNβ was tested at 200 ng/ml in the presence or absence of 10 or 100 µg/ml heparin. Bound IFNβ was detected by rabbit anti-hIFNβ (0.25 µg/ml, Peprotech) followed by goat anti-rabbit HRP (Southern Bio). 50 µl of HRP substrate solution was added for developing, and the reaction was stopped by adding 50 µl of 1 M H_2_SO_4_. The absorbance at 450 nm was measured by a plate reader.

Binding affinity of WT and mutant murine IFNβ to murine IFNAR1 (Acro Biosystems, IF1-M5225, His-tagged) was measured by coating 96-well plate with 100 ng mIFNAR1 and the binding was assessed using IFNβ (or mutants) concentrations from 100 ng to 30 µg/ml. Bound IFNβ was detected by rabbit anti-hIFNβ (1:3000, Cell Signaling #97450) followed by goat anti-rabbit HRP (Southern Bio). Apparent *K*_d_ value was calculated using Prism software.

### Disaccharide analysis of HS

BMMs from wildtype or *Rbpj^ΔM^* mice were treated with TNF for 0 or 2 days in 6-well plates. At desired end point, cells were washed with PBS and digested with 3 ml pronase E (160 µg/mL, Sigma-Aldrich) in 40 mM NaOAc, 300mM NaCl, pH6.5 at 37 °C overnight. After proteolyzed, the digestion mixture was centrifuged at 3000 rpm for 15 min, and the supernatant (which contains HS) is collected for DEAE purification. Before loading to DEAE column, 2 μL^13^C-labeled N-sulfated K5 polysaccharide (45 ng/μl) was added to supernatant. After loading, the DEAE column was washed with 1.5 ml buffer A (20 mM Tris, pH 7.5 and 50 mM NaCl), followed by elution with 1.5 ml buffer B (20 mM Tris, pH 7.5 and 1 M NaCl). The eluted solution was subjected to a 3kDa MWCO spin column and centrifuged at 15,000 rpm for 10 minutes at room temperature. The retentate was kept in the 3kDa MWCO column and washed three times with 200 μl of deionized water at 15,000 rpm for 10 minutes at room temperature to get rid of salt in the sample solution. The desalted HS was subjected to heparin lyases digestion. Samples were digested in 100 μl heparin lyases digestion solution containing 7.5 μl enzymatic buffer (100 mM sodium acetate, 2 mM calcium acetate buffer and 0.1 g/L BSA, pH 7.0), 4 μl heparin lyase I (0.5 mg/ml) and 2.5 μl heparin lyase II (13.9 mg/ml). All heparin lyases were expressed in *E.coli* and purified in house. The digestion solution was incubated at 37 °C for 12 h, after which it was boiled at 100 °C for 10 min. Before recovering the digests from the digest solution, a known amount ^13^C-labeled disaccharide calibrants (△[^13^C]UA-GlcNAc, △[^13^C]UA2S-GlcNAc, △[^13^C]UA-GlcNAc6S, △[^13^C]UA2S-GlcNAc6S, △[^13^C]UA-GlcNS, △[^13^C]UA2S-GlcNS, △[^13^C]UA-GlcNS6S and △[^13^C]UA2S-GlcNS6S) were added to the digestion solution. The HS disaccharides were recovered by centrifugation, and supernatant were freeze-dried for AMAC derivatization and LC-MS/MS analysis using our published protocol (*62*).

### Statistical analysis

Statistical analyses were performed using GraphPad Prism^®^ software. For comparisons between two groups, a two-tailed Student’s t test was used. In the case of more than two groups of samples, one-way analysis of variance (ANOVA) was used with one condition, and two-way ANOVA was used with more than two conditions. When ANOVA was performed, post hoc Bonferroni’s correction was applied for multiple comparisons. A p value < 0.05 was considered statistically significant. Data are presented as mean ± SD, as indicated in the figure legends.

### Data availability

The datasets generated during this study have been deposited in the Gene Expression Omnibus (GEO) under accession code GSE306898. Source data are provided with this manuscript.

## ACKNOWLEDGEMENTS

We thank Dr. Kazuki Inoue for his valuable contribution to the development of this project. We are grateful to Dr. Ruoxi Yuan for valuable advice on bioinformatics, Dr. Matthew Greenblatt for access to the μCT equipment, and Dr. Jeffery Esko for providing the Hs2st1 floxed mice. We are grateful to the lab members from Dr. Baohong Zhao’s laboratory for their helpful discussions and assistance. This work was supported by grants from the National Institutes of Health (AR078212 (BZ), AG093457 (BZ), AR071463 (BZ), DE031273 (DX, JL), GM142304 (ZW)) and by support for the Rosensweig Genomics Center at the Hospital for Special Surgery from The Tow Foundation. The content of this manuscript is solely the responsibilities of the authors and does not necessarily represent the official views of the NIH.

## AUTHOR CONTRIBUTIONS

T.Z. and H.H. designed and performed the experiments, analyzed and curated data, prepared figures and source files, and contributed to manuscript preparation. Specifically, T.Z. generated and finalized Fig. 1–5, Fig. 6I–O, and Supplementary Figures, and H.H. generated and finalized Fig. 1D, Fig. 2C, Fig. 6 and Supplementary Tables. J.J. performed all bioinformatic analyses and contributed to manuscript preparation. Z.W., M.L., C.N., J.L. assisted with experiments. D.X. conceived and supervised the heparan sulfate related experiments and wrote the manuscript. B.Z. conceived and directed the overall project and wrote the manuscript. All authors reviewed, provided input on the manuscript and approved submission.

## Competing Financial Interests statement

Zhangjie Wang is an employee at Glycan Therapeutics. Jian Liu is a founder of Glycan Therapeutics and has stock ownership of the company. The other authors declare that no competing interests exist.

## SUPPLEMENTAL INFORMATION

**Suppl Fig. 1 *Hs2st1* deletion does not influence RANKL-induced osteoclastogenesis nor basal/physiological bone mass.** (A) Osteoclast differentiation induced by RANKL (40 ng/ml) for 4 days determined by TRAP staining (Upper) and the number of TRAP-positive multinucleated cells per well (Bottom), n = 4/group. (B) Histomorphometry analysis of the slices of the trabecular bone. n = 9/group. (C) μCT images and bone morphometric analysis of the distal femurs isolated from the indicated 8-week-old male littermate mice. N =12 /group. BV/TV, bone volume per tissue volume; Tb.N, trabecular number; Tb.Sp, trabecular separation; Tb.Th, trabecular thickness; Oc.S/BS, osteoclast surface/bone surface; N.Oc/B.Pm, number of osteoclasts per bone perimeter. ns, not statistically significant. Data are mean ± SD. Source data are provided as a Source Data file.

**Suppl Fig. 2 GSVA scores of the Osteoclastogenic (OC)-High and OC-Low groups**. Violin plot showing the GSVA scores of the indicated cell groups.

**Suppl Fig. 3 Time course of joint swelling of inflammatory arthritis developed in *Hs2st1^ΔM^* mice and littermate controls.** (A) The photo of the hind paws. (B) Average clinical joint scores based on the total limb scores per mouse in each group. (C) Average joint swelling based on the summed wrist and ankle thickness measurements per mouse in each group. n=6/Control group; n=9/Arthritis group. Data are mean ± SD. n.s., not statistically significant. Source data are provided as a Source Data file.

**Suppl Fig. 4 The expression of *HS2ST1 in* RA samples.**Violin plot showing the expression of *HS2ST1* in healthy, early RA, and established RA groups. Each dot represents a single sample. \**p<0.05; **p < 0.01; ***p < 0.001; ****p < 0.0001*; ns: not statistically significant. Data are mean ± SD. Source data are provided as a Source Data file.

**Suppl Fig. 5 The mutant IFNβ does not influence RANKL-induced osteoclastogenesis.** (A-B) Osteoclast differentiation of WT BMMs induced by RANKL (40ng/ml) stimulation with or without WT mIFNβ, Mut1 mIFNβ or Mut2 mIFNβ variants at the indicated concentrations for 4 days. TRAP-positive multinucleated cells (MNCs) are shown in red (A). The number of TRAP-positive multinucleated cells (≥3 nuclei/cell) per well was calculated (B). n = 3/group. Scale bar, 100 μm. Mut1 mIFNβ: K48A mutant, or Mut2 mIFNβ: K48A–K37A mutant. *p<0.05; **p < 0.01; ***p < 0.001; ****p < 0.0001; ns, not statistically significant. Data representative of at least three independent experiments. Data are mean ± SD. Source data are provided as a Source Data file.

**Suppl Table 1: Quantification of HS disaccharides produced by WT or RBP-J KO BMMs with or without TNF treatment.** Quantification of HS disaccharides levels in BMMs from WT control and *Rbpj^ΔM^* mice treated with or without TNF for 2 days, as measured by LC–MS/MS (n = 3 per group).

**Suppl Table 2: IFNβ mutants binding to heparin Sepharose column.**

