## Supplementary for "An RBP-J-heparan sulfate-dependent gatekeeper directs macrophage fate between inflammatory regulation and osteoclastogenesis"

**TZ and HH contribute equally.**

****Lead Correspondence:**
Baohong Zhao, MB, DMD, PhD
Institute for Translational Medicine and Pharmacology
Icahn School of Medicine at Mount Sinai
One Gustave L. Levy Place, Box 1218
New York, NY 10029, USA


***Co-Correspondence:**
Ding Xu, PhD
Emory University School of Medicine
Emory Musculoskeletal Institute
5 Executive Park Dr. E.,
Atlanta, GA, 30329
404-251-4326 (Tel)


### SUPPLEMENTAL INFORMATION

**Suppl Table 1: Quantification of HS disaccharides produced by WT or RBP-J KO BMMs with or without TNF treatment.** Quantification of HS disaccharides levels in BMMs from WT control and $Rbpj^{\Delta M}$ mice treated with or without TNF for 2 days, as measured by LC–MS/MS (n = 3 per group).

**Suppl Table 2: IFNβ mutants binding to heparin Sepharose column.**

**Suppl Fig. 1**

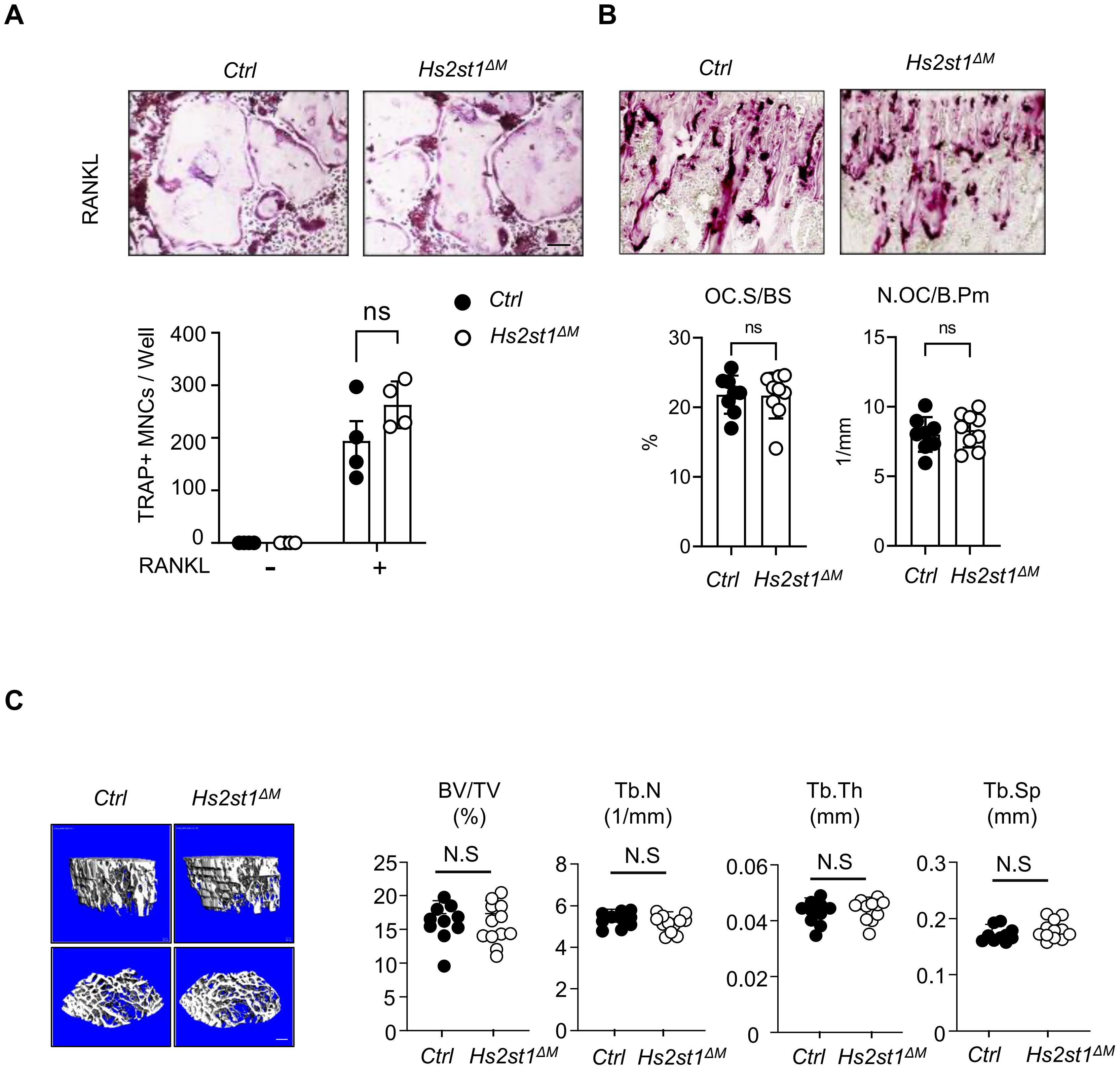
A
B
Ctrl
Hs2st1ΔM
RANKL
ns
TRAP+ MNCs / Well
400
300
200
100
0
RANKL
-
+
OC.S/BS
N.OC/B.Pm
%
1/mm
30
20
10
15
5
C
BV/TV
(%)
Tb.N
(1/mm)
Tb.Th
(mm)
Tb.Sp
(mm)
N.S
25
8
6
4
2
0.06
0.04
0.02
0.3
0.2
0.1

**Suppl Fig. 2**

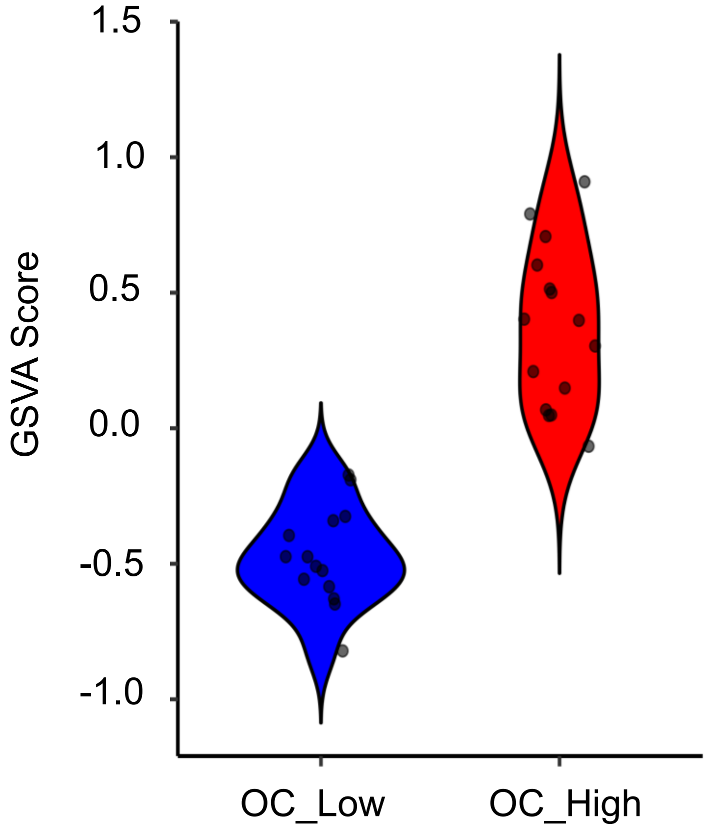

**Suppl Fig. 3**

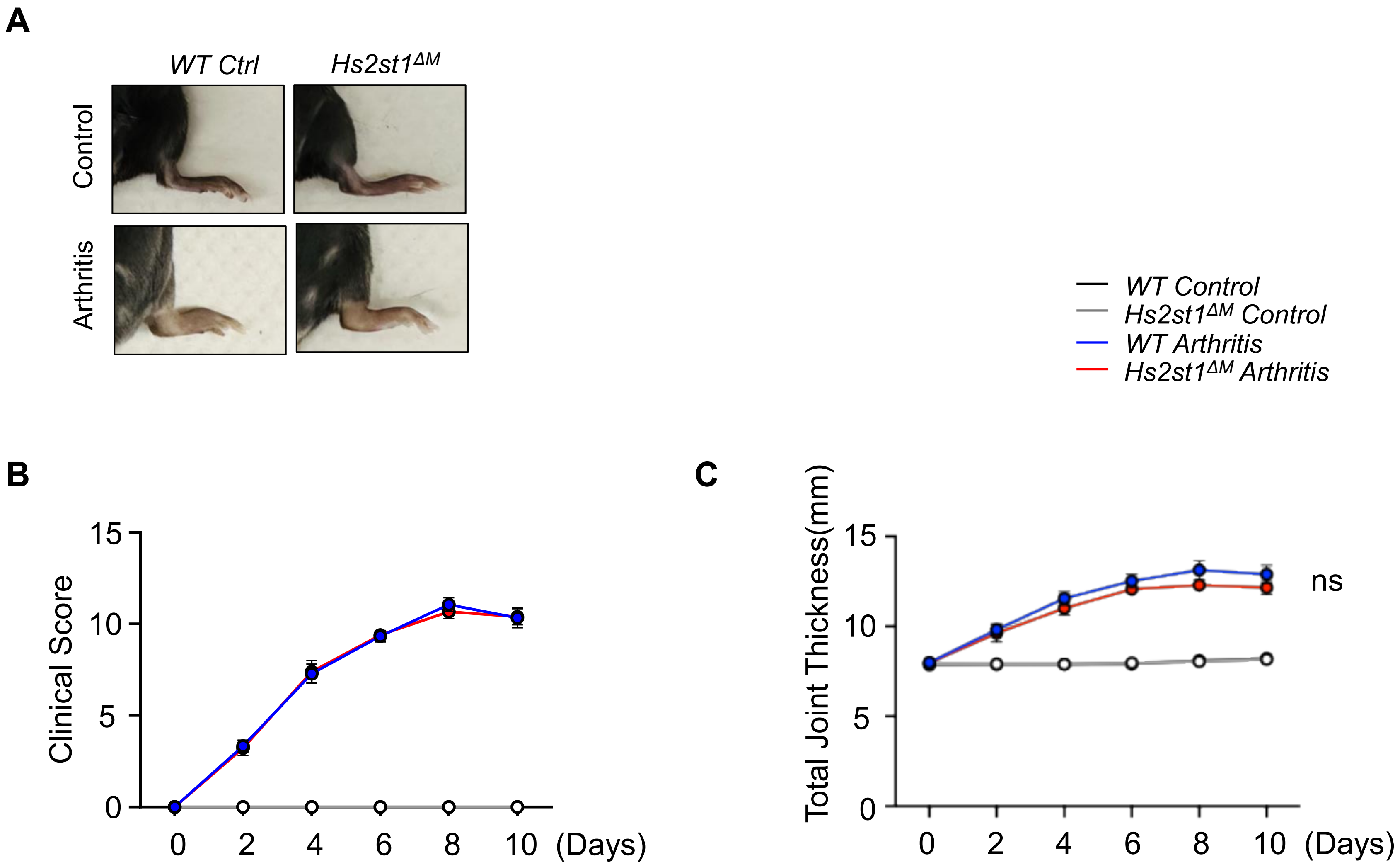

A
WT Ctrl
Hs2st1ΔM
Control
Arthritis
WT Control
Hs2st1ΔM Control
WT Arthritis
Hs2st1ΔM Arthritis
B
Clinical Score
15
10
5
0
0 2 4 6 8 10 (Days)
C
Total Joint Thickness(mm)
15
10
5
0
ns
0 2 4 6 8 10 (Days)

**Suppl Fig. 4**

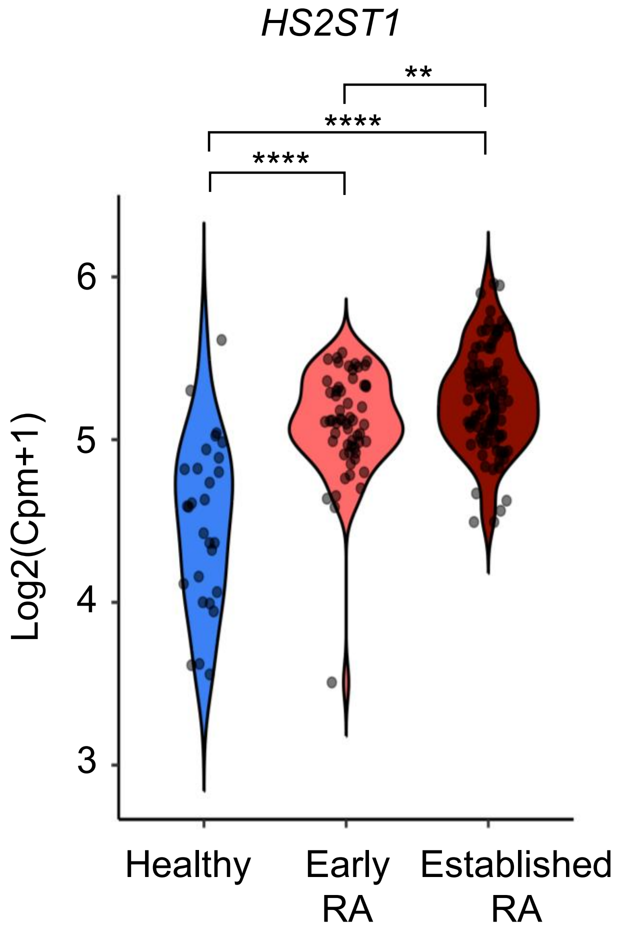

**Suppl Fig. 5**

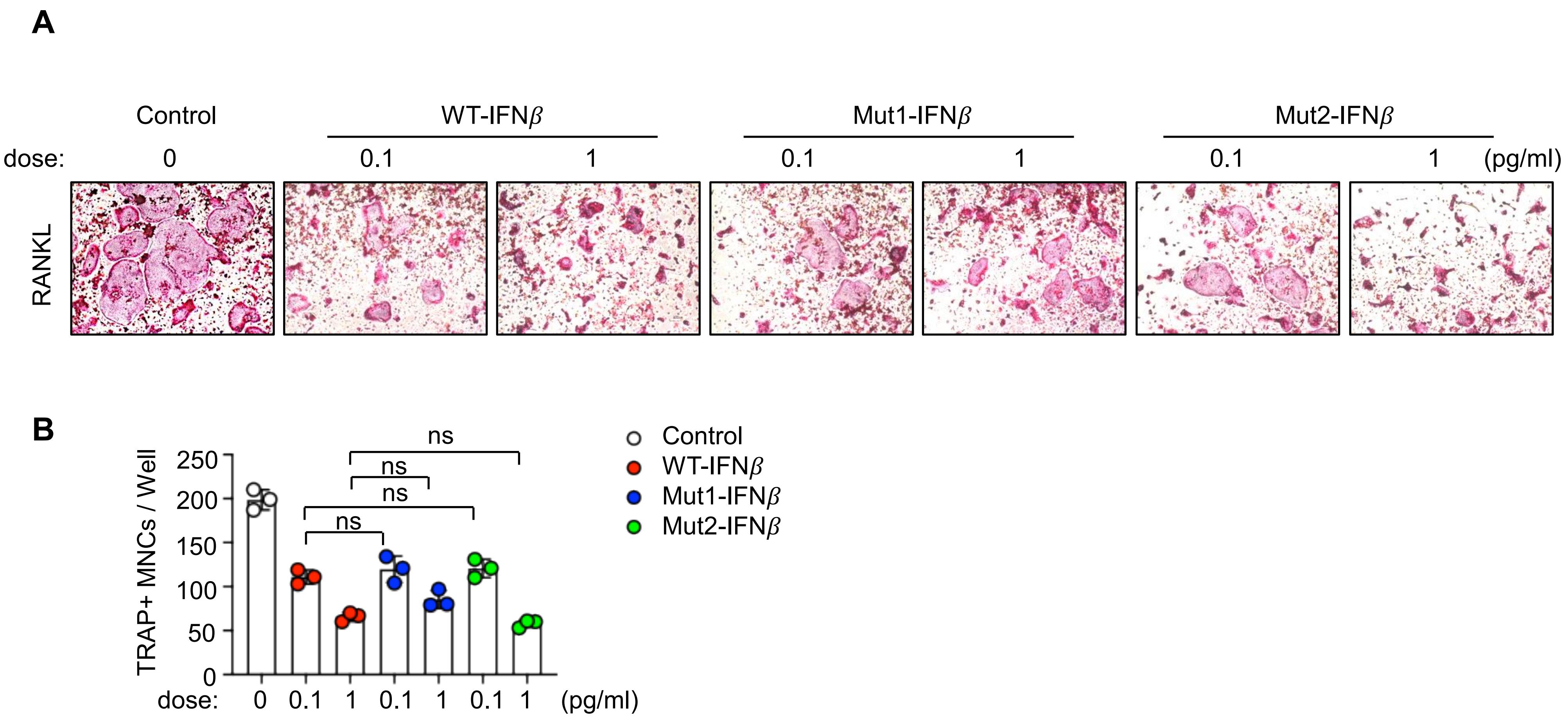
A
Control
WT-IFNβ
Mut1-IFNβ
Mut2-IFNβ
dose: 0 0.1 1 0.1 1 0.1 1 (pg/ml)
RANKL
B
TRAP+ MNCs / Well
250
200
150
100
50
0
ns
ns
ns
ns
dose: 0 0.1 1 0.1 1 0.1 1 (pg/ml)
Control
WT-IFNβ
Mut1-IFNβ
Mut2-IFNβ

Supplementary Table 1: Quantification of HS disaccharides produced by WT or RBP-J KO BMMs with or without TNF treatment

| Disaccharides | ng | | | | | | | | | | | |
|---|---|---|---|---|---|---|---|---|---|---|---|---|
| | **WT** | **WT** | **WT** | **KO** | **KO** | **KO** | **WT TNF** | **WT TNF** | **WT TNF** | **KO TNF** | **KO TNF** | **KO TNF** |
| △UA2S-GlcNS6S | 4.37 | 3.94 | 3.98 | 4.02 | 5.04 | 2.49 | 11.17 | 9.83 | 7.11 | 11.59 | 9.55 | 10.09 |
| △UA-GlcNS6S | 0.96 | 1.70 | 2.12 | 2.51 | 1.96 | 1.81 | 4.52 | 3.23 | 2.17 | 3.95 | 4.17 | 3.56 |
| △UA2S-GlcNS | 3.54 | 5.24 | 4.29 | 6.10 | 7.47 | 5.50 | 12.54 | 8.90 | 6.62 | 14.83 | 14.84 | 14.18 |
| △UA-GlcNS | 6.98 | 7.18 | 7.51 | 8.91 | 7.73 | 7.65 | 12.47 | 11.96 | 9.82 | 21.95 | 21.01 | 20.46 |
| △UA2S-GlcNAc6S | 0.38 | 0.54 | 0.48 | 0.55 | 0.63 | 0.59 | 0.68 | 0.71 | 0.52 | 0.72 | 0.64 | 0.54 |
| △UA-GlcNAc6S | 2.31 | 2.31 | 2.18 | 2.95 | 2.05 | 1.77 | 3.05 | 2.87 | 1.53 | 4.49 | 4.35 | 4.09 |
| △UA2S-GlcNAc | 0.00 | 0.00 | 0.03 | 0.29 | 0.30 | 0.00 | 0.75 | 0.84 | 0.46 | 3.31 | 3.28 | 3.04 |
| △UA-GlcNAc | 1.66 | 3.28 | 3.23 | 5.15 | 4.13 | 3.45 | 9.31 | 9.06 | 6.50 | 17.58 | 16.24 | 15.85 |
| Tota HS amount | 20.19 | 24.19 | 23.83 | 30.49 | 29.31 | 23.27 | 54.49 | 47.41 | 34.72 | 78.43 | 74.07 | 71.82 |
| Total 2S containing disaccharide | 8.29 | 9.72 | 8.78 | 10.96 | 13.44 | 8.58 | 25.14 | 20.28 | 14.71 | 30.45 | 28.31 | 27.85 |
| Total 6S containing disaccharide | 8.02 | 8.49 | 8.76 | 10.03 | 9.68 | 6.66 | 19.42 | 16.64 | 11.33 | 20.75 | 18.71 | 18.28 |
| Total NS containing disaccharide | 15.85 | 18.06 | 17.9 | 21.54 | 22.2 | 17.45 | 40.7 | 33.92 | 25.72 | 52.32 | 49.57 | 48.29 |

Supplementary Table 2: IFNβ mutants binding to heparin Sepharose column

| Mutant | Elution NaCl concentration(mM) |
|---|---|
| WT | 585 |
| K37A | 510 |
| R123A | 520 |
| R126A | 510 |
| K129A | 560 |
| K132A | 560 |
| K45A | 490 |
| K48A | 470 |
| K101A | 490 |
| R105A | 510 |
| K37A-R123A | 490 |
| K48A-K101A | 465 |
| K48A-K37A | 435 |
| K48A-R123A | 450 |
